# Nb*PTR1* confers resistance against *Pseudomonas syringae pv. actinidiae* in kiwifruit

**DOI:** 10.1101/2023.09.07.556601

**Authors:** Shin-Mei Yeh, Minsoo Yoon, Sidney Scott, Abhishek Chatterjee, Lauren M. Hemara, Ronan K.Y. Chen, Tianchi Wang, Kerry Templeton, Erik H.A. Rikkerink, Jay Jayaraman, Cyril Brendolise

**Affiliations:** The New Zealand Institute for Plant & Food Research Limited (PFR), Mt Albert Research Centre, Auckland 1025, New Zealand; School of Biological Sciences, University of Auckland, Auckland 1010, New Zealand; The New Zealand Institute for Plant and Food Research Limited (PFR), Palmerston North 4474, New Zealand; The New Zealand Institute for Plant and Food Research Limited (PFR), Motueka 7198, New Zealand

**Keywords:** HopZ5, Effector-trigged immunity, RNAi hairpin library, *Nicotiana benthamiana*, Psa3, *Actinidia chinensis*

## Abstract

*Pseudomonas syringae pv. actinidiae* biovar 3 (Psa3) causes a devastating canker disease in yellow-fleshed kiwifruit (*Actinidia chinensis*). The effector HopZ5, which is present in all isolates of Psa3 causing global outbreaks of pandemic kiwifruit canker disease, triggers immunity in *Nicotiana benthamiana* and is not recognised in susceptible *A. chinensis* cultivars. In a search for *N. benthamiana* non-host resistance genes against HopZ5, we found that the nucleotide-binding leucine-rich repeat receptor NbPTR1 recognised HopZ5. RPM1-interacting protein 4 (RIN4) orthologues from multiple plants, including kiwifruit, were associated with NbPTR1-mediated autoimmunity suppression and recognition of HopZ5. No functional orthologues of Nb*PTR1* were found in *A. chinensis*. Nb*PTR1* transformed into Psa3-susceptible *A. chinensis* var. *chinensis* ‘Hort16A’ plants introduced HopZ5-specific resistance against Psa3. Altogether, this study suggested that expressing NbPTR1 in Psa3-susceptible kiwifruit is a viable approach to acquiring resistance to Psa3 and it provides valuable information for engineering resistance in otherwise susceptible kiwifruit genotypes.

## Introduction

*Pseudomonas* syringae pv. *actinidiae* (Psa) is a bacterial plant pathogen which causes a devastating canker disease in kiwifruit. Symptoms of Psa infection include blossom necrosis, cane dieback, trunk/cane bleeding, cankerous growths, and necrotic leaf spots ^1^. Psa was first isolated in Japan (in the 1980s) with subsequent outbreaks reported throughout China, Korea, Italy and New Zealand ^2–5^. Since 2008, the emergence of a highly virulent lineage of Psa biovar 3 (Psa3; also called the pandemic lineage of Psa3) has led to significant losses in kiwifruit production worldwide. This pandemic lineage is particularly virulent towards yellow-fleshed kiwifruit (*Actinidia chinensis*), which is a major crop for both Italy and New Zealand ^4,5^. Currently, Psa’s impact is mainly mitigated through hygienic orchard practices. However, the development of more tolerant cultivars will play a key role in the long-term management of the disease. Achieving robust Psa resistance has therefore become an important target of kiwifruit breeding programmes.

Plants have evolved two key modes of defence against invading pathogens such as Psa. The first is pattern-triggered immunity (PTI), which relies upon the recognition of pathogen-associated molecular patterns (PAMPs) by specific pattern-recognition receptors (PRRs) at the plant cell-surface ^6^. Pathogenic bacteria have in turn developed the ability to overcome PTI by injecting host cells with virulence proteins called effectors. Effectors are delivered into host cells through a type III secretion system (T3SS) and can facilitate infection by interfering with pattern recognition and allowing pathogens to evade immune detection ^7^. To counteract effectors, plants have developed a second layer of defence called effector-triggered immunity (ETI), which involves intracellular receptors to recognise bacterial effectors and triggers the immune response. ETI produces a robust immune response by restoring and potentiating PTI ^8^. ETI often results in a form of programmed cell death known as the hypersensitive response (HR), which allows plants to kill off infected cells to prevent further disease spread ^9,10^.

ETI is governed by the aforementioned intracellular, nucleotide-binding domain, leucine-rich repeat receptors commonly referred to as NLRs ^10^. NLRs can sense effectors that are secreted by pathogens into plant cells by a variety of mechanisms, including a type III secretion system (T3SS) carried by some bacterial pathogens like Psa. Effectors translocated into plant cells via the T3SS are known as type III effectors (T3Es) ^11^ and are commonly named Hop (Hrp outer protein) or Avr (avirulence) proteins ^12^. Psa3 carries approximately 39 T3Es, including two effectors (HopZ5 and HopH1) that are unique to this highly virulent biovar ^13^. Several *P. syringae* T3E proteins target host RPM1-INTERACTING PROTEIN 4 (RIN4) but trigger ETI owing to the NLRs guarding RIN4 ^14^. The *P. syringae* pv. *tomato* effector AvrRpt2 cleaves RIN4 to generate a product involved in activating RPS2-mediated immunity in *Arabidopsis thaliana* ^15,16^. Similar RIN4 cleavage mechanisms have been found for Mr5-mediated defence responses in apple ^17^ and Ptr1-mediated recognition of AvrRpt2 in tomato ^18^. In soybean, RIN4 modification is involved in the recognition of AvrB and AvrRpm1 by Rpg1b and Rpg1r, respectively ^19^. Recently, Jayaraman et al. ^20^found that five pathogenicity-associated Psa biovar 3 effectors (HopZ5a, HopH1a, AvrPto1b, AvrRpm1a, and HopF1e) are collectively essential for full Psa3 virulence and that they largely target host RIN4 proteins. Although most Psa T3Es have been identified, we have yet to fully uncover which NLR proteins are responsible for recognizing and responding to specific effectors during ETI. Previously, Brendolise et al. ^21^ constructed a hairpin-RNAi library targeting 345 NLR gene candidates identified in *Nicotiana benthamiana* ^21^. This library has been used to identify several NLR genes that are able to recognize effectors from Psa. For example, NRG1 (N requirement gene 1) and Rpa1 (Resistance to *Pseudomonas syringae* pv. *actinidiae* 1) have been shown to be involved in the recognition of Psa effectors HopQ1 and AvrRpm1, respectively ^22,23^.

The Psa effector HopZ5 is a member of the *Yersinia* outer protein J (YopJ) effector family and has an acetyltransferase activity that triggers HR in *A. thaliana* and *N. benthamiana* ^24,25^. Several R genes have been shown to mediate HopZ5-triggered immunity in these model species. *RPM1* (*RESISTANCE TO P. SYRINGAE PV. MACULICOLA* 1) is required for HopZ5-triggered immunity in *Arabidopsis*, responding to HopZ5’s acetylation of threonine residue T166 of AtRIN4 ^26^. In *N. benthamiana,* HopZ5 recognition depends on several NLR genes, *PTR1* (*PSEUDOMONAS TOMATO RACE 1*) and *ZAR1* (*HOPZ-ACTIVATED RESISTANCE 1*) with the assistance of *JIM2* (*XOPJ4 IMMUNITY 2*) for the latter ^27^. The evaluation of a set of Psa3 effector knockouts suggests that HopZ5 is not recognised in susceptible *A. chinensis* ^28^. Hence, we hypothesized that introducing HopZ5-recognizing *R* genes into *A. chinensis* could be a promising approach for facilitating ETI and improving resistance to Psa. This study demonstrates the independent identification of *PTR1a* as an NLR which recognizes HopZ5 in *N. benthamiana*, as well as the transformation of *NbPTR1a* into previously susceptible *A. chinensis*. We show that Nb*PTR1a*-expressing *A. chinensis* var. *chinensis* ‘Hort16A’ plants display significantly improved resistance to Psa3 via reduced *in planta* bacterial growth and disease symptoms, owing to a specific HopZ5-triggered resistance response.

## Results

### NbPTR1a mediates HopZ5-triggered immunity in *Nicotiana benthamiana*

Previous studies have demonstrated that HopZ5 triggers a significant hypersensitive response (HR) in *Nicotiana* spp.^24^. To identify the NLR gene(s) responsible for mediating HopZ5 recognition in *Nicotiana benthamiana*, HopZ5 was screened against a library of NLR gene-silencing RNAi hairpin constructs following the methodology previously described ^21^. One hairpin construct (hp#1) was shown through transient expression assays to interfere with HopZ5-triggered HR (Figure 1a). hp#1 contains six DNA fragments, each targeting different NLR genes (with some occasional overlap): m1, m2, u10, u120, u147, and u156 ^21^. To identify the fragment responsible for interfering with HopZ5-triggered HR, each hairpin fragment was individually sub-cloned and screened. Only fragment m1 was able to reduce the HopZ5-triggered HR (Figure 1b). m1 targets two genes within the *N. benthamiana* genome ^29^: Nb*PTR1a* (NbS00012936g0019.1) and Nb*PTR1b* (NbS00007796g0005.1) (Figure 1c). Nb*PTR1b* is rendered non-functional owing to the presence of a premature stop codon, suggesting that Nb*PTR1a* is likely to encode the NLR required for this immune response.

**Figure 1.**
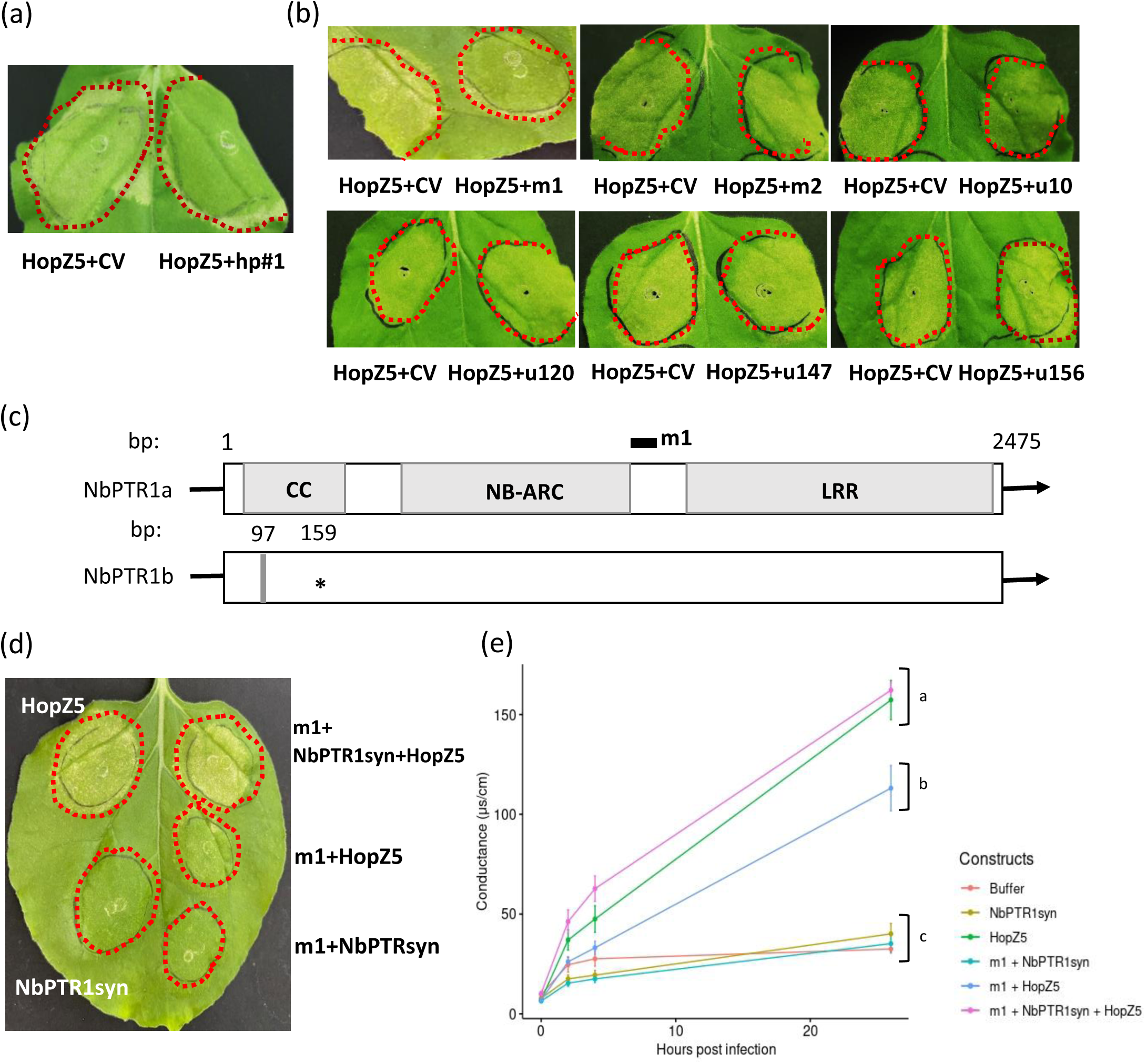
NbPTR1a mediates a HopZ5-triggered immune response in *Nicotiana benthamiana*. **a,b** HopZ5-triggered HR is reduced by hp#1 and the m1 fragment. *N. benthamiana* leaves were agroinfiltrated with pTKO2_GGT control vector (CV), hp#1, or one of six DNA target fragments in hp#1: m1, m2, u10, u120, u147, u156, each at OD_600_ of 0.2, followed by agroinfiltration of HopZ5 (OD_600_ of 0.05) 48 h later. Leaves were photographed 6 days post-inoculation (dpi). Red dash line indicated the HopZ5-inoculated area. **c** Schematic of PTR1 gene structure in *N. benthamiana* (Nb*PTR1a* and Nb*PTR1b*). The coiled-coil (CC), nucleotide binding site present in APAF-1, R proteins and CED-4 (NB-ARC), and leucine-rich repeat (LRR) domains are indicated in grey rectangles. The black bar indicates the area targeted by m1, corresponding to position 1291-1441bp relative to Nb*PTR1a*. Nb*PTR1b* has a 5-bp deletion (grey bar) at position 97 bp resulting in a pre-mature stop codon (asterisk) at position 159 bp. **d** Transient complementation of Nb*PTR1syn*. *N. benthamiana* leaves were agroinfiltrated with m1+Nb*PTR1syn*, m1+GUS, CV+Nb*PTR1syn* or CV+GUS, each at OD_600_ of 0.1, followed by agroinfiltration with HopZ5 or GUS control (each at OD_600_ of 0.2) 48 h later. Leaves were photographed 4 dpi. **e** Quantification of the hypersensitive response (HR) by electrolyte leakage. Conductivity was measured from leaf disks collected at 2 day post final infiltration from the leaf patches shown in (**d**). Error bars represent the standard errors of the means for ten independent biological replicates, collected from two independent experimental runs (n=10). HopZ5 was used as positive control and infiltration buffer (10mM MgCl_2_, 100 μM acetosyringone) as a negative control. Letters indicate statistically significant differences from a one-way ANOVA and Tukey’s HSD *post hoc* test for values at 26 h post sampling.

To confirm that NbPTR1a is required for HopZ5 triggered immunity, endogenous Nb*PTR1a* was silenced and complemented with a synthetic analogue Nb*PTR1syn*. NbPTR1syn amino acid sequence is identical to NbPTR1a but has a modified nucleotide sequence to elude silencing by the m1 hairpin construct (Figure S1). Transient co-expression of Nb*PTR1syn* with HopZ5 was able to restore HR to the Nb*PTR1a*-silenced leaf patches, supporting the idea that functional Nb*PTR1a* is required for the recognition of HopZ5 (Figure 1d). HR was also quantified by measuring the conductivity due to ion leakage from the corresponding infiltrated patches. Co-expression of m1 and Nb*PTR1syn* in the presence of HopZ5 showed a significant restoration of ion leakage (Figure 1e). These results indicated that NbPTR1a could recognize HopZ5 to trigger HR in *N. benthamiana,* consistent with the previous observations by Ahn et al.^27^.

While HopZ5-triggered HR was reduced by m1 expression in the transient assay, ion leakage was only partially reduced (Figure 1e). Ahn *et al.* ^27^ found that Nb*ZAR1* was also involved in the recognition of HopZ5 in *N. benthamiana.* Therefore, we identified two silencing fragments from our hairpin library, m16 and u38, which correspond to the nucleotide-binding-ARC (NB-ARC) domain or the coiled-coil (CC) domain of Nb*ZAR1*, respectively (Figure S2a). Neither silencing construct alone nor the combined m16/u38 constructs were able to block HopZ5-triggered HR to the extent that silencing Nb*PTR1a* could (Figure S2b and S2c). To check if silencing constructs were efficiently silencing their targets, endogenous Nb*PTR1* and Nb*ZAR1* expression was checked. RT-qPCR of Nb*PTR1* and Nb*ZAR1* showed that Nb*PTR1* expression is reduced by *PTR1*-targeting m1 but not with *ZAR1*-targeting m16/u38; whereas Nb*ZAR1* expression is reduced with m16/u38 but not with m1, as expected (Figure S2d). These collective results did not identify Nb*ZAR1* as a significant contributor to HopZ5-triggered HR under these experimental conditions.

### RIN4 is involved in the recognition of HopZ5 in *N. benthamiana*

*Ptr1* from *Solanum lycopersicoides* can cause autoimmunity when overexpressed in *Nicotiana glutinosa* leaves, and this autoimmunity response is suppressed by co-expression of three tomato *RIN4* genes ^30^. Yoon & Rikkerink ^23^ have previously cloned multiple *RIN4* orthologues from *N. benthamiana*, *A. chinensis* var. *chinensis* ‘Hort16A’ and *Arabidopsis thaliana* Col-0. To assess whether these RIN4 orthologues can suppress Nb*PTR1a* autoimmunity when overexpressed in *N. benthamiana*, Nb*PTR1a* and *RIN4* orthologues were delivered simultaneously or sequentially into *N. benthamiana* leaves and the reduction of Nb*PTR1a* autoimmunity response (reduced cell death) was tested. NbPTR1a-mediated autoimmunity was suppressed by three Nb*RIN4*s (Nb*RIN4-1*, Nb*RIN4-2*, and Nb*RIN4-3*) and two AcRIN4s (Ac*RIN4-2* and Ac*RIN4-3*) when the *RIN4* constructs were infiltrated 48 h earlier than Nb*PTR1a* (Figure S3). This result suggests that NbRIN4-1, NbRIN4-2, NbRIN4-3, AcRIN4-2, and AcRIN4-3 may be involved in suppressing NbPTR1a activity, possibly through a direct interaction with NbPTR1a.

Physical association of NbPTR1a with the RIN4 orthologues were tested by co-immunoprecipitation following the procedures described previously by Yoon and Rikkerink ^23^. Interestingly, of the three NbRIN4 orthologues, only NbRIN4-1 was able to interact stably with NbPTR1a, despite all three NbRIN4 orthologues being able to suppress NbPTR1a-triggered autoimmunity (Figure S4). We then tested the ability of NbPTR1a to interact with AtRIN4 and the three kiwifruit RIN4 homologues (Figure 2a). Surprisingly, only AtRIN4 and AcRIN4-2 showed strong physical interactions with NbPTR1a, while a weak or no interaction was found with AcRIN4-1 and AcRIN4-3, respectively.

**Figure 2.**
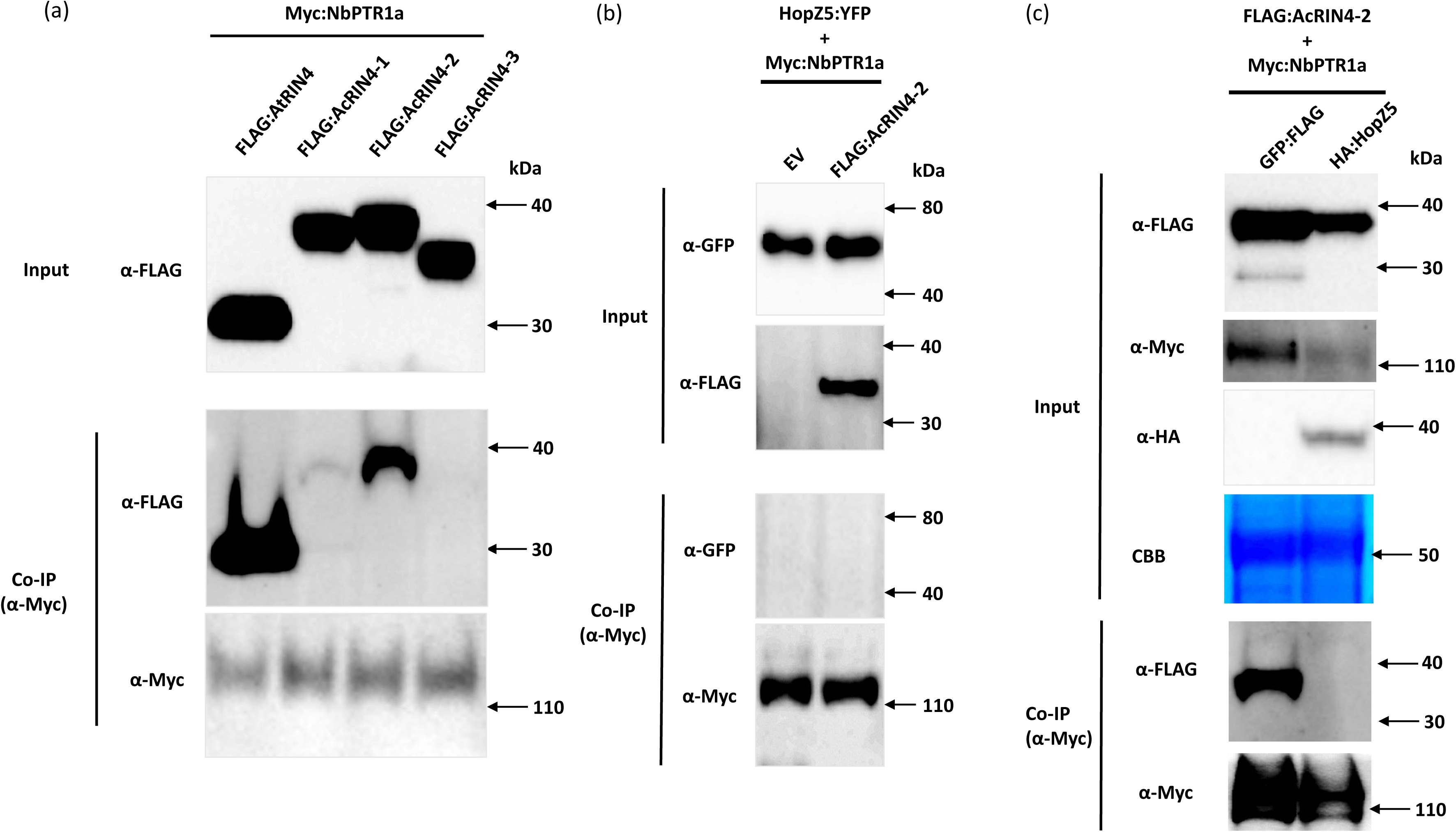
AcRIN4-2 is involved in NbPTR1a-mediated recognition of HopZ5. **a** Co-immunoprecipitation of NbPTR1a and RIN4 homologues. NbPTR1a and RIN4 homologues were co-expressed simultaneously via agroinfiltration, each at OD_600_ of 0.1. 2 days post-infiltration, leaf samples were harvested, protein extracts prepared and precipitated using anti-Myc antibody. Western blot of input and precipitated proteins were probed with anti-FLAG or anti-Myc antibody. The experiments were conducted twice with similar results. **b** Co-expression of AcRIN4-2 with NbPTR1a does not facilitate HopZ5a immunoprecipitation with NbPTR1a. YFP-tagged HopZ5 (or YFP alone), Myc-tagged NbPTR1a, and FLAG-tagged AcRIN4-2 were expressed simultaneously by agroinfiltration, each at OD_600_ of 0.1. 2 days post-infiltration, leaf samples were harvested, and protein extracts prepared and precipitated using anti-Myc antibody. Western blots of input and precipitated proteins were probed with anti-GFP or anti-Myc antibody. IP, co-immunoprecipitation. **c** HopZ5 abolishes co-immunoprecipitation of AcRIN4-2 with NbPTR1a. YFP-tagged HopZ5 (or YFP alone), Myc-tagged NbPTR1a, and FLAG-tagged AcRIN4-2 were expressed simultaneously by agroinfiltration, each at OD_600_ of 0.1. 2 days post-infiltration, leaf samples were harvested, and protein extracts prepared and precipitated using anti-Myc antibody. Western blots of input and precipitated proteins were probed with anti-GFP, anti-FLAG, or anti-Myc antibody. IP, co-immunoprecipitation; CBB, coomassie brilliant blue.

Recently, Jayaraman et al. ^20^ found that HopZ5 interacted with AcRIN4-1 and AcRIN4-2 and weakly with AcRIN4-3. To determine the potential association between HopZ5 and NbPTR1a with a bridging interaction by AcRIN4-2, we performed a three-component co-immunoprecipitation experiment with co-expression of NbPTR1a:Myc, HopZ5:YFP, and FLAG:AcRIN4-2 (Figure 2b). We found that in when NbPTR1a was immunoprecipitated in the presence of AcRIN4-2, HopZ5 was not co-precipitated. These results suggested either, that despite strongly interacting with HopZ5 and NbPTR1a, AcRIN4-2 does not bridge the HopZ5-PTR1a interaction, or that HopZ5 modifies AcRIN4-2 in a way that eliminates interaction with NbPTR1a. To assess this possibility, AcRIN4-2 was checked for interaction with NbPTR1a, in the presence and absence of HopZ5 (Figure 2c). Notably, the presence/activity of HopZ5 abolished the ability of AcRIN4-2 to interact with NbPTR1a, probably allowing NbPTR1a to trigger immunity as a consequence.

### Ac*PTR1* homologues could not complement loss of Nb*PTR1a* in *N. benthamiana*

To assess if any functional homologues (orthologues) of Nb*PTR1a* were present in Psa3-susceptible *A. chinensis* var. *chinensis*, the recently published ‘Red5’ genome was searched for Nb*PTR1a* homologues ^31^. Two kiwifruit *PTR1a* homologues were identified in ‘Red5’ and genes amplified from *A. chinensis* var. *chinensis* ‘Hort16A’ genomic DNA. Both alleles for each gene were cloned: Ac*PTR1a*-1, Ac*PTR1a*-2, Ac*PTR1b*-1, and Ac*PTR1b*-2 (Figure S5). The nucleotide sequence of Nb*PTR1a* is 55% and 56% identical to those of Ac*PTR1a* and Ac*PTR1b*, respectively. To confirm whether the four Ac*PTR1*s were the closest kiwifruit homologues of Nb*PTR1a*, a reciprocal BLAST search was conducted for each of the Ac*PTR1* homologues’ NB-ARC domains against the *N. benthamiana* genome. Four gene candidates were identified: NbS00012936 (Nb*PTR1a*), NbS00007796 (Nb*PTR1b*), NbS00034734, and NbS00047736. A phylogenetic analysis of the NB-ARC domain of the *N. benthamiana* candidates and Ac*PTR1* homologues showed that the four Ac*PTR1* homologues formed a single group phylogenetically closer to the clade with NbS00034734 and NbS00047736 than that of Nb*PTR1a* (Figure 3a). However, the lengths of the two *N. benthamiana* genes are much shorter than those of the Ac*PTR1* homologues or Nb*PTR1a*; NbS00034734 has a truncated NB-ARC domain and NbS00047736 lacks its LRR domain. This result suggested that, while not likely to be the closest true homologues of Nb*PTR1a*, the identified Ac*PTR1a*/Ac*PTR1b* candidates are probably the closest functional NLRs to Nb*PTR1a* present in ‘Hort16A’.

**Figure 3.**
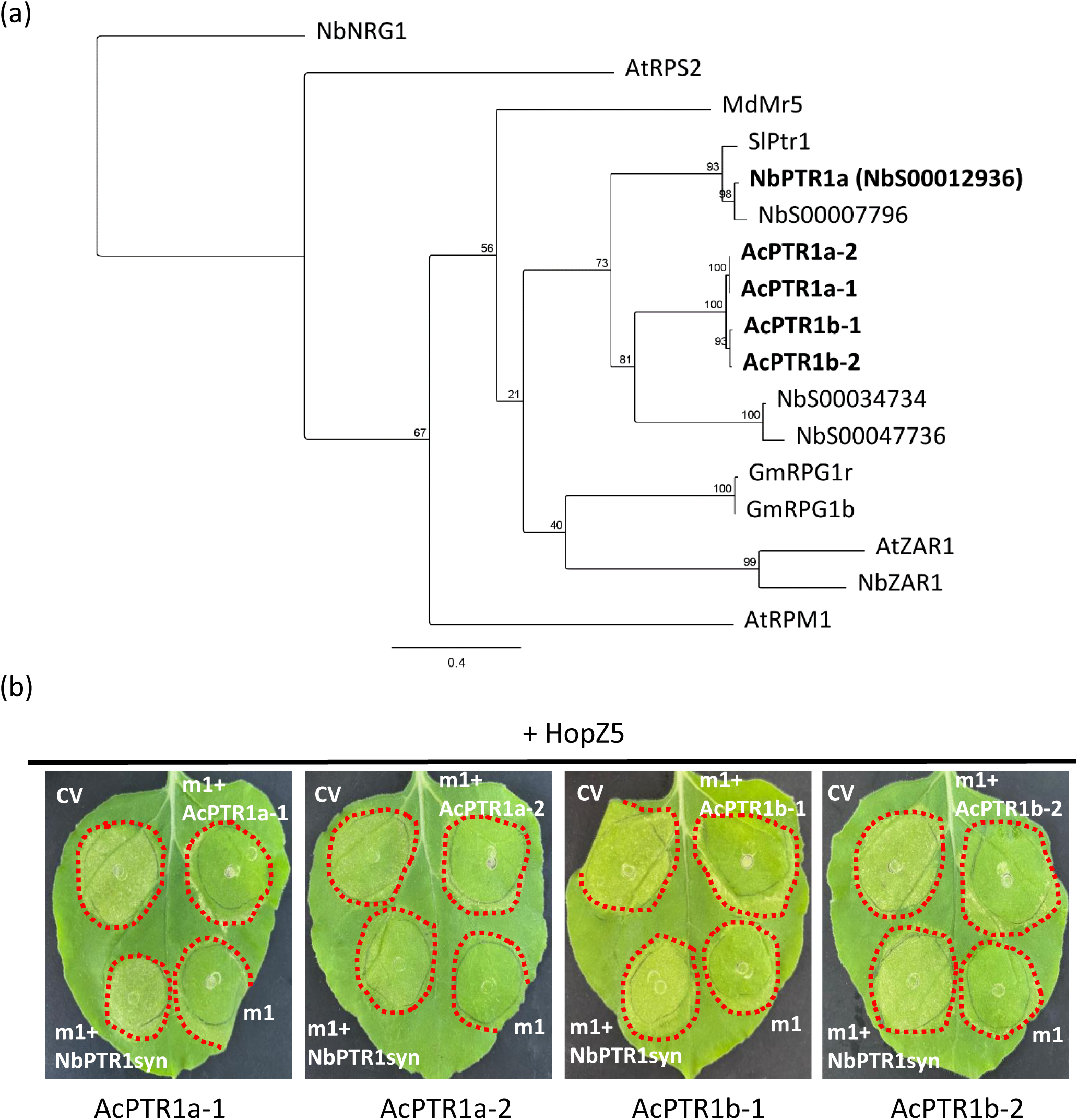
Ac*PTR1* homologues cannot complement silencing of Nb*PTR1a* in *Nicotiana benthamiana*. **a** Phylogenetic relationship of R genes involved in the recognition of RIN4-associated effectors including Ac*PTR1* homologues and their most closely related genes in *N. benthamiana*. Nucleotide sequences of NB-ARC domains were aligned by using MUSCLE alignment. The nucleotide distance was calculated by GTR GAMMA model and the tree was constructed by using RAxML with 100 bootstrap replicates in Geneious Prime. Sequences were sourced from *N. benthamiana* (Nb), *Solanum lycopersicoides* (Sl), *Arabidopsis thaliana* (At), *Actinidia chinensis* (Ac), *Glycine max* (Gm), or *Malus domestica* (Md). The tree was drawn to scale. Labels on branches indicate the percentage of bootstrap support with Nb*NRG1* used as outgroup. R genes of interest are indicated in bold font. **b** Transient complementation of kiwifruit *PTR1a* homologues. Four *PTR1a* homologues were identified in kiwifruit: Ac*PTR1a-1*, Ac*PTR1a-2*, Ac*PTR1b-1*, and Ac*PTR1b-2*. *N. benthamiana* leaves were agroinfiltrated with the m1 hairpin construct and individual Ac*PTR1* homologues, as indicated, each at OD_600_ of 0.05, followed by agroinfiltration with HopZ5 (OD_600_ of 0.2) 48 h later. Leaves were photographed 4 dpi. Red dashed lines indicate the HopZ5-inoculated area. The experiments were performed in at least three independent leaves, across three independent experiments with similar results. CV: control vector.

Several RIN4-related NLR genes have been identified, including At*RPS2*, Md*Mr5*, Sl*Ptr1*, Gm*RPG1r*, Gm*RPG1b*, At*RPM1*, At*ZAR1*, and Nb*ZAR1* ^15,17–19,26,27^. To understand the broader relationships of Ac*PTR1* homologues with these NLR proteins involved in recognition of RIN4-associated effectors, the NB-ARC domains of each NLR protein were also included in our phylogenetic analysis (Figure 3a). This revealed that that there is strong support for the clade shared by Ac*PTR1* homologues and Nb*PTR1a*/Sl*Ptr1*, but weak support for exclusion of the clade with Gm*RPG1b*/Gm*RPG1r* and At*ZAR1*/Nb*ZAR1*, suggesting a RIN4-guarding function for the AcPTR1 homologues.

To assess whether the four identified kiwifruit *PTR1* homologues could recognise HopZ5 in *N. benthamiana*, endogenous Nb*PTR1a* was silenced by the m1 construct and complemented by each Ac*PTR1* homologue (Figure 3b). None of the Ac*PTR1* homologues was able to restore HopZ5-triggered HR. These results indicate that the four identified kiwifruit PTR1 homologues do not function like NbPTR1a to recognize HopZ5 in *N. benthamiana*, suggesting that no PTR1-like orthologue exists in ‘Hort16A’ plants, commensurate with their Psa3-susceptible status.

### Overexpression of Nb*PTR1a* in kiwifruit confers resistance against Psa3 infection

Hemara *et al.* ^28^ previously found that HopZ5 was not recognised in susceptible *A. chinensis* var. *chinensis* ‘Hort16A’ plants and did not trigger an HR. Both RPM1 and PTR1 play an important role in HopZ5-triggered immunity in *A. thaliana* and *N. benthamiana*, respectively ^26,27^. This suggested that transformation of either At*RPM1* or Nb*PTR1* into kiwifruit might trigger recognition of HopZ5 and be associated with resistance to Psa3. Therefore, stable ‘Hort16A’ transformants expressing either At*RPM1* or Nb*PTR1a* under the control of a 35S CaMV promoter were generated. The resulting transgenic plantlets (grown in axenic tissue culture) were flood-inoculated with Psa3 ICMP 18884 (also called Psa3 V-13). Plantlets were monitored for development of classic bacterial canker symptoms over a 6-week period to assess resistance. Similarly to β-glucuronidase (GUS) control transgenic plants, most At*RPM1* transgenic plantlets did not survive for six weeks post-infection, whereas the majority of Nb*PTR1a* transgenic plantlets showed moderate to high survival under similar inoculation conditions (Table S1). These preliminary results suggested that the Nb*PTR1a* transgenic ‘Hort16A’ plants are more resistant to Psa3 than At*RPM1* transgenic ‘Hort16A’ plants.

Amongst the Nb*PTR1a* transgenic plants, Line 1A was consistently susceptible to Psa3 infection (Table S1). The level of expression of the Nb*PTR1a* transgene was quantified by qPCR and revealed that the five selected Nb*PTR1a* transgenic lines showed varied levels of Nb*PTR1a* expression (Figure 4a). Interestingly, Line 1A showed a very low level of expression of the transgene, consistent with a lack of resistance to Psa3 measured in our preliminary experiments. Furthermore, no differences in the growth phenotypes between the NbPTR1a transgenic plants and wild-type Hort16A plants. To further assess the resistance to Psa3 of the Nb*PTR1a* transgenic plants, an *in planta* bacterial growth assay was conducted on mature glasshouse plants. Fully expanded leaves were inoculated and Psa3 growth was quantified at seven days post-inoculation (dpi) by PDQeX-qPCR (Figure 4b). At 7 dpi the four moderate-to-high expressing lines (4, 7D, 13A, and 14) significantly restricted *in planta* Psa3 growth, whereas the very low expressing Line 1A showed no significant difference from wild-type ‘Hort16A’ or GUS control plants. These results showed that the expression of the Nb*PTR1a* transgene conferred resistance to Psa3 in ‘Hort16A’ in both immature tissue culture and mature glasshouse plants.

**Figure 4.**
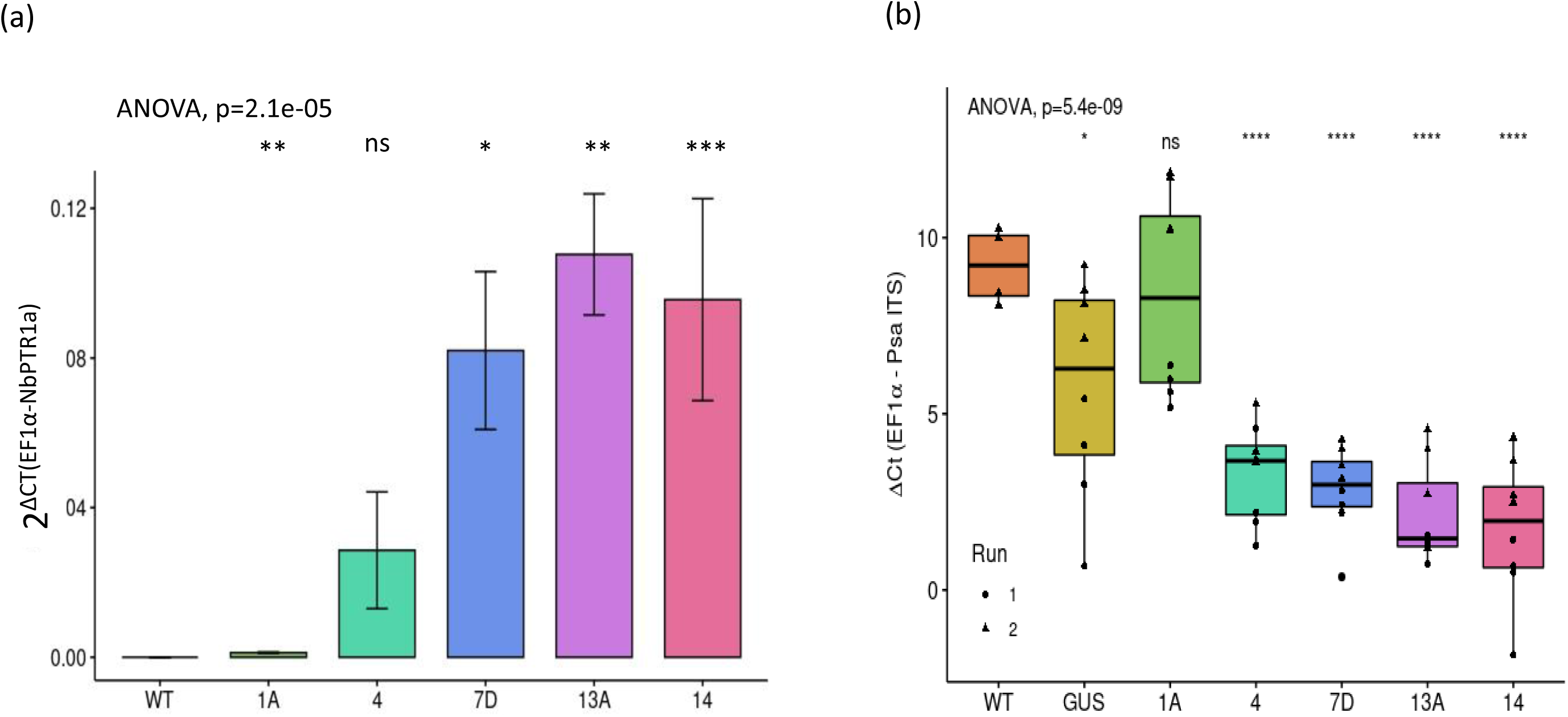
Nb*PTR1a* transgenic kiwifruit plants are resistant to Psa3. **a** qPCR quantification of *PTR1* expression in five selected Nb*PTR1a* transgenic lines. The Nb*PTR1a* transgene expression is normalised to reference gene, Ac*EF1α*. Wild-type Hort16A plants were used as a negative control. Data were shown as means ± SEM of three biological replicates. Asterisks indicate results of a one-way analysis of variance (ANOVA) and a two-tailed Welch’s *post-hoc* t-test between the selected transgenic lines and wild-type Hort16A; (*) P < 0.05, (**) P < 0.01, (***) P < 0.001, and (ns) non-significant. **b** Bacterial growth quantification in Nb*PTR1a* transgenic glasshouse-grown plants. Leaves were inoculated with Psa3 at approximately 10^7^ CFU/mL and bacterial growth was determined at seven days post infection (dpi). Wild-type Hort16A and β-glucuronidase (GUS) transgenic lines were used as negative controls. Error bars represent standard error of the mean from four biological replicates. Asterisks indicate results of a one-way analysis of variance (ANOVA) and a two-tailed Welch’s *post-hoc* t-test between the selected transgenic lines and wild-type Hort16A; (*) P < 0.05, (**) P < 0.01, (***) P < 0.001, (****) P < 0.0001, and (ns) non-significant.

To further confirm that NbPTR1a-mediated growth restriction was specifically related to the recognition of HopZ5, the five selected transgenic lines were grown in tissue culture and flood-inoculated with wild-type Psa3 or Psa3 Δ*hopZ5*, a Psa3 strain mutated to lack the *hopZ5* effector gene ^28^. The disease phenotypes from flood-inoculated plantlets were assessed for up to 6 weeks post-inoculation. The disease symptoms included necrosis/leaf wilting, leaf spots, and white spots with numerical categories assigned (Figure S6a). Disease phenotypes were quantified using an adapted ‘area under the disease progression curve’ (AUDPC) methodology (Figure S6b)^32^. The disease symptomology indicated that the four Nb*PTR1a* moderate-to-high expressing lines (4, 7D, 13A, and 14) showed a significant decrease in symptom development over the 6 weeks when inoculated with Psa3, whereas no difference in symptom development for these lines was observed in comparison to the GUS control and low expression Line 1A plants when plantlets were inoculated with Psa3 Δ*hopZ5* (Figure 5a). Next, the plantlets were inoculated under the same conditions with Psa3 or Psa3 Δ*hopZ5* strains and the *in planta* bacterial growth was quantified at ten dpi by PDQeX-qPCR (Figure 5b). Similarly to the disease progression results, *in planta* growth of Psa3 was greatly restricted for in the four moderate-to-high expressing lines (4, 7D, 13A, and 14) compared with the GUS control and low-expressing Line 1A plantlets. No similar growth restriction was measured with the Psa3 Δ*hopZ5* strain, suggesting that Psa3 growth restriction is specifically due to HopZ5 recognition. Additionally, the disease symptomology and *in planta* growth assays of the tissue culture-grown Nb*PTR1a* plantlets reflected closely the results obtained previously with the glasshouse-grown plants.

**Figure 5.**
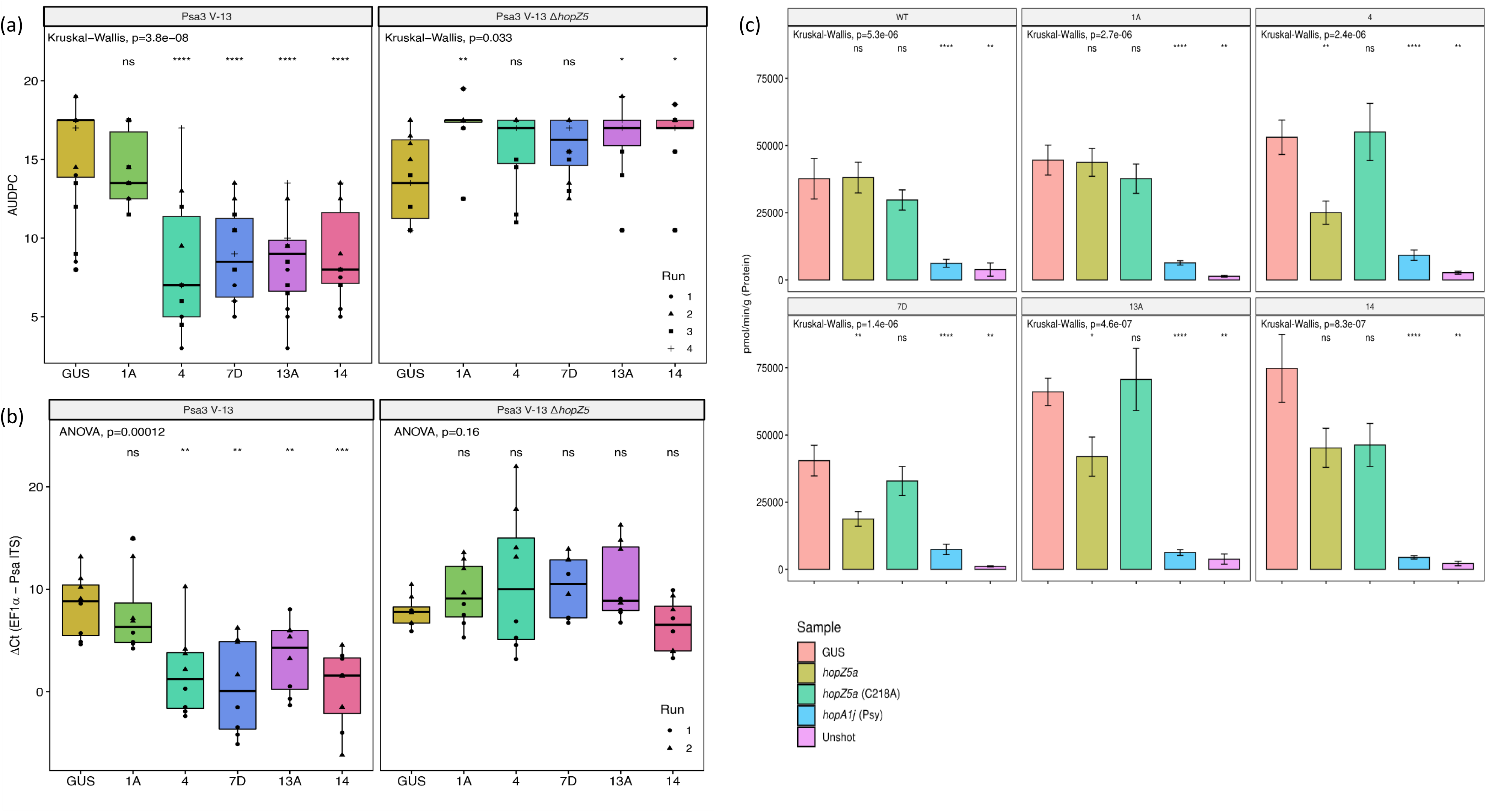
PTR1a-triggered bacterial growth restriction *in planta* is HopZ5-specific. **a** Disease phenotyping analysis of Nb*PTR1a* transgenic plantlets inoculated with Psa3 V-13 or Psa3 V-13 *ΔhopZ5*. Plantlets were flood-inoculated with Psa3 V-13 or Psa3 V-13 *ΔhopZ5* at approximately 10^6^ CFU/mL. Individual plantlets were scored weekly on a scale of 0 (asymptomatic) to 4 (dead) for 6 weeks post-infection (dpi). Disease phenotypes were quantified by measuring the area under the disease progression curve (AUDPC). Asterisks indicate results of a Kruskal-Wallis analysis of variance and a two-tailed Welch’s *post-hoc* t-test between the selected transgenic lines and transgenic GUS control; (*) P < 0.05, (**) P < 0.01, (***) P < 0.001, (****) P < 0.0001, and (ns) non-significant. Data were collected across four independent experimental runs. **b** Bacterial growth in Nb*PTR1a* transgenic plantlets determined by PDQeX-qPCR. Psa3 V-13 or Psa3 V-13 *ΔhopZ5* was flood-inoculated as described in (**a**), and bacterial presence in leaf samples was quantified at 10 days post-inoculation by PDQeX DNA extraction and qPCR for the Psa3 intergenic transcribed spacer (*ITS*) region normalized to Ac*EF1α*. Error bars represent standard error of the mean from four biological replicates. Asterisks indicate results of a one-way analysis of variance (ANOVA) and a two-tailed Welch’s *post-hoc* t-test between the selected transgenic lines and transgenic GUS control line; (*) P < 0.05, (**) P < 0.01, (***) P < 0.001, (****) P < 0.0001, and (ns) non-significant. Data from two independent experimental runs are displayed. **c** HopZ5 triggers immunity in Nb*PTR1a* transgenic plants. HopZ5-triggered HR was monitored using a reporter eclipse assay. Effector constructs tagged with GFP, or empty vector, were co-expressed with a GUS reporter construct using biolistic bombardment in leaves from five Nb*PTR1a* transgenic lines or wild-type Hort16A. GUS activity was measured 48 h after DNA bombardment. Error bars represent the standard errors of the means for three independent biological replicates with six technical replicates each (n = 18). HopA1j from *Pseudomonas syringae* pv. *syringae* 61 was used as positive control and un-infiltrated leaf tissue (Unshot) as a negative control. Asterisks indicate results of a Kruskal-Wallis analysis of variance and a Wilcoxon *post-hoc* test between the selected effector treatments and empty vector (GUS) alone; (*) P < 0.05, (**) P < 0.01, (***) P < 0.001, (****) P < 0.0001, and (ns) non-significant. Data were collected across three independent experimental runs.

To assess whether the Nb*PTR1a* transgenic plants could specifically recognize HopZ5 activity to trigger resistance to Psa3, a biolistic transformation reporter eclipse assay was conducted on tissue culture plantlet leaves as previously described by Jayaraman *et al.* ^33^. A GUS reporter gene was co-expressed together with either HopZ5 or its non-functional HopZ5-C218A mutant by DNA bombardment in the leaves of the transgenic lines (Figure 5c). The effector HopA1 from *P. syringae* pv. *syringae* 61 was used as a positive control for HR in this assay ^33^. Three transgenic lines (4, 7D, and 13A) showed significant reduction in GUS activity when co-bombarded with HopZ5 but not with the enzymatically dead HopZ5-C218A in comparison to the GUS bombardment alone. These results indicated that Nb*PTR1a* transgenic plants could specifically recognize HopZ5 acetyltransferase activity to trigger an HR. There was a small, non-significant decrease in GUS activity visible when comparing GUS bombardment alone with HopZ5 for Line 14 but this was surprisingly no different from the response to HopZ5-C218A.

To confirm that NbPTR1a mediates the recognition of HopZ5 in transgenic plants, transgenic glasshouse plant leaves were vacuum-infiltrated with Psa3, Psa3 Δ*hopZ5*, or Psa3 Δ*hrcC,* and HR was evaluated by ion leakage measurement (Figure S7). The Psa3 Δ*hrcC* strain was used as a negative control for HR in this assay owing to its lack of a type III secretion system and an associated inability to cause an effector-triggered HR. Surprisingly, no significant difference for HR-associated ion leakage was visible in these Nb*PTR1a* transgenic plants when inoculated with Psa3 or Psa3 Δ*hopZ5,* similar to that observed in GUS control plants. Psa3 Δ*hrcC* showed low leakage as expected owing to its inability to secrete effectors. Due to the ability of Psa3 to suppress effector-triggered HR likely though effector-effector interference^33^, *P. fluorescens* PF0-1 artificially carrying the type III secretion system from *P. syringae* pv. *syringae*, called Pfo(T3S)^34^ as used to deliver HopZ5 into the transgenic NbPTR1 and GUS plants (Figure S8). The enzymatically dead HopZ5-C218A was used as negative control^25^ and HopA1j from *P. syringae* pv. *syringae* 61 used as positive control ^33^. HopZ5 appeared to trigger some ion leakage in the four moderate-to-high expressing lines (4, 7D, 13A, and 14) compared with the GUS control and low-expressing Line 1A plantlets. Taken together, these results indicate that HopZ5 was able to trigger a weak but detectible HR in Nb*PTR1a* transgenic plants under our experimental conditions.

## Discussion

In this study, using an RNAi hairpin screening library we demonstrated that NbPTR1a recognised Psa3 effector HopZ5 in *N. benthamiana*. Furthermore, we identified that no functional kiwifruit *PTR1* orthologues capable of recognising HopZ5 are present in Psa3-susceptible ‘Hort16A’ kiwifruit plants. We also showed that RIN4 is probably involved with Nb*PTR1a*-mediated recognition of HopZ5, either directly through interaction with PTR1 or through a third interactor. Finally, we transformed Nb*PTR1a* into ‘Hort16A’ to confer the first transgenic resistance against Psa3 introduced into a commercial kiwifruit cultivar. Overall, we present that a PTR1-like function is not present in kiwifruit and required complementation by Nb*PTR1a* for resistance to Psa3.

Previously, two nucleotide-binding leucine-rich repeat proteins (NLR) in *N. benthamiana* were identified using an NbNLR VIGS screening library as responsible for mediating HopZ5 recognition: *PTR1* and *ZAR1* (the latter with the assistance of *JIM2*) ^27,35^. From our RNAi hairpin library screening, however, we identified a role only for *PTR1* and not for *ZAR1* in the recognition of HopZ5 in *N. benthamiana*. Two silencing fragments, m16 and u38, both corresponding to Nb*ZAR1,* were unable to reduce HopZ5-triggered HR in *N. benthamiana* (Figure S2). While the possibility remains that the two Nb*ZAR1*-targting fragments could not fully silence Nb*ZAR1*, it is clear that silencing Nb*PTR1* alone was sufficient to eliminate HopZ5-triggered HR. Meanwhile, Zheng *et al.* ^35^ found that HopZ5 triggered HR in wild-type *N. benthamiana* but not in the *zar1* mutant, without PTR1 involvement. Although it is not clear why this discrepancy was observed, *N. benthamiana* plants used for *Agrobacterium*-mediated transient expression by Ahn *et al.* (NbPTR1 and NbZAR1) and Zheng *et al*. (NbZAR1 only) could be genetically different from those used in our study (NbPTR1 only).

The mechanisms of several plant NLR proteins in model plant species that recognize HopZ5 have been studied. In *Arabidopsis,* RPM1, responsible for Pto DC3000-delivered HopZ5-triggered immunity, is activated by the acetylation of RIN4 by HopZ5 ^26^. ZAR1 orthologues from *A. thaliana* and *N. benthamiana* recognise HopZ5 by interacting with ZED1 and JIM2, respectively ^26,27^. While the mechanisms of At*RPM1*-, At*ZAR1*- and Nb*ZAR1*-mediated recognition of HopZ5 are clear, little is known about the mechanism for Nb*PTR1*-mediated recognition of HopZ5. Sl*PTR1* from *Solanum lycopersicoides* is required for recognition of AvrRpt2 and RipBN effectors when co-expressed with SlRIN4-3, which also supressed SlPTR1 autoimmunity in *N. glutinosa* transient expression experiments ^30^. This accumulated evidence suggested that Nb*PTR1*-mediated recognition of HopZ5 might be related to RIN4.

Previously, Mazo-Molina *et al.* ^30^ found that Sl*Ptr1* autoimmunity-associated cell death was suppressed by three tomato RIN4 proteins in *N. glutinosa* leaves. In our study, a similar autoimmunity response was also observed from over-expression of NbPTR1a in *N. benthamiana* leaves and three NbRIN4s, AcRIN4-2 and AcRIN4-3 could suppress NbPTR1a-mediated cell death (Figure S3). Interestingly, suppression of NbPTR1a autoimmunity by these RIN4s was observed only when the RIN4s were expressed 48 h earlier than NbPTR1a but not when NbPTR1a and RIN4 were expressed simultaneously. Co-immunoprecipitation (co-IP) experiments showed that NbPTR1a stably interacted with NbRIN4-1 and AcRIN4-2 (both PTR1a autoimmunity suppressors), as well as AtRIN4 (a non-suppressor of PTR1a autoimmunity). This suggested that factors beyond a stable interaction with PTR1 may be necessary for autoimmunity suppression and activation of resistance, perhaps often through a transient interaction process or another RIN4/PTR1 interacting partner, such as Exo70 ^36^. AcRIN4-2 is also targeted by HopZ5 ^20^. However, a three-way HopZ5:AcRIN4-2:NbPTR1a complex was not detected in our experiments. This surprising result could be explained in several ways: firstly, the interaction between NbPTR1a and HopZ5 (particularly due to HopZ5 activity) may be too transient; secondly, the interaction may link to unknown mechanisms and protein partners or multi-protein complexes during HopZ5-triggered immunity in *N. benthamiana*; or finally, there may be interference in the system to detect the interaction. Yeast has previously been utilized as a heterologous model system to functionally characterise T3E proteins based on its low redundancy of eukaryotic processes and lack of immunity mechanisms that counteract effectors to mask effector function ^37^. It will be interesting to test in future work whether the interaction of the HopZ5:AcRIN4-2:NbPTR1a complex could be detected using a yeast three-hybrid system. Jayaraman *et al.* ^20^ found that four out of five key redundant Psa3 effectors collectively essential for full Psa3 virulence probably targeted host RIN4 proteins (HopZ5a, AvrPto1b, AvrRpm1a, and HopF1e). Since only HopZ5 appears to be responsible for Nb*PTR1a*-conferred immunity against Psa3, this is probably further evidence that RIN4 itself may not be the host protein directly guarded by PTR1. Identifying other HopZ5 plant targets in kiwifruit or *N. benthamiana* may reveal this PTR1 guardee in the future.

Introducing *NbPTR1a* into kiwifruit facilitated recognition of HopZ5 and conferred resistance to Psa3 infection in susceptible ‘Hort16A’ plants. Among the five selected Nb*PTR1a* transgenic lines, the strength of Psa3 resistance may not be proportional to the Nb*PTR1a* gene expression level, but require a certain minimum threshold expression level to facilitate Psa3 resistance. The very low-expressing transgenic Line 1A, like the GUS control plants, was not able to recognise HopZ5. The other four moderate-to-high *NbPTR1a* expressing transgenic lines (4, 7D, 13A, and 14) showed similar levels of Psa3 resistance from disease phenotyping analysis, *in planta* bacterial growth quantification, or biolistic transformation reporter eclipse assays (Figs 4 and 5). Psa3 resistance in Nb*PTR1a* transgenic kiwifruit is stably maintained from young tissue-cultured plantlets through to mature glasshouse-grown plants. Unlike Nb*PTR1a* transgenic kiwifruit, introducing At*RPM1* into kiwifruit did not improve resistance against Psa3 infection (Table S1). Choi *et al.* ^26^ demonstrated that At*RPM1*-mediated HopZ5 recognition in *Arabidopsis* requires alteration of residue T166 in RIN4. It is possible that this recognition relies on co-evolved guardee and resistance proteins and the key modification may thus be restricted to *Arabidopsis* alleles of RIN4. Perhaps co-transforming At*RIN4* with At*RPM1* into kiwifruit may facilitate the recognition of HopZ5 to improve Psa3 resistance in the At*RPM1* transgenic plants. Alternatively, an RPM1-like function may already exist in ‘Hort16A’ but is somehow overcome by the effector complement present in Psa3 in a manner that does not affect NbPTR1a signalling. As AtRPM1 guards RIN4, such a suppression would constitute further evidence that RIN4 modifications may not be directly responsible for NbPTR1a signalling. Evidence for this lies in recognition of AvrRpm1_Pma_ in ‘Hort16A’ plants when delivered by biolistic assay or *Pseudomonas fluorescens* (T3S), but not bacterial growth assay of Psa3 carrying AvrRpm1_Pma_ ^33^.

We show that PTR1a recognition of Psa3 is HopZ5-specific in the kiwifruit transgenic plants. HopZ5 is able to trigger a weak HR in these Nb*PTR1a* transgenic plants (Fig. S8), but only in isolation from other Psa3 effectors (Fig. S7), similar to what was previously observed in Psa3-resistant *Actinidia arguta* AA07_03 ^28^. It will be interesting to study whether a *PTR1* homolog exists in *A. arguta* and can confer resistance to Psa3. The lack of ion leakage in response to HopZ5 recognition in both *Actinidia* species when delivered by Psa3 is unexpected, and the molecular mechanism underlying the Nb*PTR1a*-mediated recognition of HopZ5 remains unclear.

We have discussed previously how breeding durable resistance genes into targeted kiwifruit cultivars will play an important role in long-term management of Psa ^28^. The introduction of a functional *PTR1* gene in kiwifruit through crosses with resistant plant lines carrying a functional kiwifruit orthologue of *PTR1* will allow it to be efficiently tracked as it is backcrossed into various breeding lines. However, traditional plant breeding can be time-consuming and slow the development of new varieties ^38^. Alternatively, the new method of CRISPR/Cas-based ‘Prime editing’ has been widely applied in precision plant breeding era ^38^. Recently, this GE technology has been used in kiwifruit by deleting flowering regulation genes to reduce plant dormancy ^39^. Such a strategy, but with a lighter-touch editing approach, might be possible to ‘repair’ any non-functional PTR1 orthologues present in kiwifruit. This would potentially be a more efficient way to introduce resistance genes without the linkage drag caused by undesirable traits in more classical breeding approaches used to develop Psa-resistant kiwifruit varieties.

## Materials and methods

### Bacterial strains

*Agrobacterium tumefaciens* strain GV3101 was used for Agrobacterium-mediated transient assay (agroinfiltration) in *N. benthamiana,* and *A. tumefaciens* strain EHA105 was used for kiwifruit transformation ^40^. *Agrobacterium* strains were grown on lysogeny broth (LB) with appropriate antibiotics at 28°C. *E. coli* DH5α was used for plasmid maintenance and grown in LB medium at 37°C. All strains were stored in 20% glycerol+ LB at −80°C.

Knockout strains of *P. syringae* pv. *actinidiae* ICMP 18884 biovar 3 (Psa3 V-13) *ΔhrcC* and *ΔhopZ5* were previously generated through transposon mutagenesis and homologous recombination, respectively ^28,41^. The Psa3 Δ*hrcC* strain contains all Psa3 effectors but without a functional type III secretion system to delivery effector proteins into a host plant ^33^. Bacteria were streaked from glycerol stocks onto LB agar supplemented with 12.5 μg/mL nitrofurantoin and 40 μg/mL cephalexin (Sigma Aldrich).

### Cloning and gene synthesis

Nb*PTR1* genes were identified by BLAST from *N. benthamiana* (version 0.4.4) ^29^ and PCR products were amplified from *N. benthamiana* genomic DNA. Kiwifruit *PTR1* candidate homologues were identified by BLAST from the *A. chinensis* var. *chinensis* ‘Red5’ genome ^31^. Kiwifruit *PTR1* candidate homologues were PCR amplified from *A. chinensis* var. *chinensis* ‘Hort16A’ genomic DNA. Sequences of all the primers are provided in Table S2.

PCR products of candidate genes were purified using the Zymoclean Gel DNA Recovery kit (Zymo Research) and then cloned into pENTR/D/SD entry vector using the In-Fusion cloning kit (Takara). Inserts of all plasmids were sequenced by Macrogen, Korea. The silencing fragment amplified from hairpin constructs were cloned in the pTKO2 hairpin vector ^21^. The glycosyltransferases gene, *F3GGT1 (GGT),* from red-fleshed kiwifruit (*Actinidia chinensis*) ^42^ was used as the control fragment for hairpin silencing fragments and the pENTR-GGT entry clone was recombined by Gateway cloning into pTKO_2_ ^21^ to generate the pTKO_2_-GGT control vector. The full-length candidates were cloned in the pHEX2 vector using Gateway cloning (ThermoFisher Scientific), respectively. Similarly, the control vector pHEX2-GUS was constructed by Gateway LR reaction of the pENTR-GUS (Invitrogen, USA) into pHEX2 vector ^43^.

The nucleotide sequence of synthetic Nb*PTR1* (NbPTR1syn) was designed using the GenSmart™ Codon Optimization tool (https://www.genscript.com/tools/gensmart-codon-optimization) with manual editing (Fig. S2). NbPTR1syn was synthesized by GenScript (Singapore) and cloned into the pHEX2 vector using Gateway cloning (ThermoFisher Scientific, USA).

### RNA/DNA extraction and qPCR

Genomic DNA from *A. chinensis* var. *chinensis* ‘Hort16A’, *A. thaliana* Col-0, or *N. benthamiana* leaf was extracted using the DNeasy Plant Mini Kit (Qiagen), following the manufacturer’s instructions. RNA from young kiwifruit tissue culture leaves was extracted using the Spectrum Plant Total RNA kit (Sigma-Aldrich), following the manufacturer’s protocol except the incubation temperature of lysed tissue sample was increased to 65°C.

To check the expression of Nb*PTR1a* in transgenic plants, cDNA was synthesised using extracted RNA as the template and the QuantiTect Reverse Transcription kit (Qiagen) by following the manufacturer’s instructions. qPCR was performed with specific primers (Table S2) and LightCycler 480 SYBR Green I Mastermix (Roche) using the LightCycler 480 II real-time PCR device (Roche). Each reaction volume was 10 µL and reactions were run in quadruplicate, including non-template negative controls. qPCR followed a three-step reaction of 95°C for 10 s, 60°C for 10 s and 72°C for 20 s for 50 cycles. The data were analysed by the LightCycler 480 software 1.5 using the target/reference ratio to compare the target gene expression level to that of the reference gene, AcEF1α. Sequences of all primers are provided in Table S2.

### Plant materials and growth condition

For transient expression experiments, *N. benthamiana* plants were grown in a temperature-controlled glasshouse at 22 to 24°C under long day conditions (16 h light, 8 h dark).

Tissue-cultured *Actinidia chinensis* var. *chinensis* ‘Hort16A’ plantlets were purchased from Multiflora Laboratories (New Zealand). Plantlets were grown on supplied Murashige and Skoog (MS) agar (M1, Table S3) in 400 mL-lidded plastic tubs. Plantlets were grown in a tissue culture room at 20°C under long day conditions (16 h light, 8 h dark) with cool white fluorescent light (∼120 μmol^−2^ m s^−1^). For glasshouse experiments, tissue culture plantlets were transferred to soil and grown for at least 3 months prior to testing. Rooted kiwifruit plants grown in the soil were fertilized with commercial WUXAL (2mL/L) every two weeks.

### Transient over-expression and RNAi library screen in *Nicotiana benthamiana*

Transient expression and the RNAi library screen in *N. benthamiana* were performed as previously described ^21^. In brief, freshly grown *Agrobacterium tumefaciens* culture containing either RNAi hairpin constructs or pTKO2_GGT control vector was re-suspended in infiltration buffer (10 mM MgCl_2_, 100 μM acetosyringone). The cells were diluted to the appropriate concentrations and infiltrated in leaves of 3-week-old plants using a needleless syringe and the infiltrated area was marked. After 48-h infiltration, the marked infiltrated area were inoculated with the HopZ5 construct or the GUS control. Photographs were taken 4–5 days after HopZ5 infiltration.

### Electrolyte leakage quantification in *N. benthamiana*

Electrolyte leakage experiments in *N. benthamiana* were performed as previously described ^27^. In brief, six leaf disks (6 mm) from the agroinfiltrated patches were collected 2 days after the last infiltration and washed in distilled water for 30 min. Leaf discs were placed in 2 mL of sterile water and the conductivity was measured over 26 h by using a LAQUAtwin EC-33 conductivity meter (Horiba, Japan). HopZ5 was used as the positive control and infiltration buffer (10mM MgCl_2_, 100 μM acetosyringone) as a negative control. The standard errors of the means were calculated from five biological replicates. Data were collected across two independent experimental runs and data from both runs were plotted (n = 10).

### Transient expression in *N. benthamiana* and co-immunoprecipitation

The transient expression in *N. benthamiana* and co-immunoprecipitation assay was described previously ^28^. Briefly, *A. tumefaciens* AGL1 (YFP-tagged effectors; (Choi *et al.*, 2017)) or GV3101 pMP90 (FLAG tagged AcRIN4s; ^23^) was freshly grown in LB with appropriate antibiotics at 28°C with shaking at 200 rpm. Cells were pelleted by centrifugation at 4000 g for 10 min and resuspended in infiltration buffer (10 mM MgCl_2_, 5 mM EGTA, 100 μM acetosyringone). Cell suspensions were diluted to a final OD_600_ of 0.1 and infiltrated into at least two fully expanded leaves of 4-to 5-week-old *N. benthamiana* plants using a needleless syringe. All *Agrobacterium*-mediated transformation experiments were performed using pre-mixed *Agrobacterium* cultures for the stipulated effector-RIN4 combinations in a single injection for co-immunoprecipitation experiments (see below). YFP was used as a negative control for effectors. Tissues (0.5 g per sample) were collected 2 days post-infiltration and ground to a homogeneous powder in liquid nitrogen and resuspended in 1 mL of protein extraction buffer (1× PBS, 1% n-dodecyl β-d-maltoside or DDM (Invitrogen, Carlsbad, CA, USA), and 0.1 tablet cOmplete™ protease inhibitor cocktail (Sigma-Aldrich) in NativePAGE™ buffer (Invitrogen). Extracted protein samples were centrifuged at 20,000 *g* for 2 min at 4°C and the supernatant was collected for immunoprecipitation using the Pierce™ Anti-c-Myc Magnetic Beads (88843; Thermo-Fisher Scientific) according to the manufacturer’s instructions. Total and immunoprecipitated proteins were resolved on a 4–12% SDS-PAGE gel. Western blots using PVDF membranes were prepared and probed using HRP-conjugated antibodies in 0.2% I-Block (Invitrogen). Detection was achieved using the Clarity™ MAX Western ECL Substrate (Bio-Rad, Hercules, CA). The antibodies used were α-FLAG-HRP (A8592; Sigma-Aldrich), α-GFP-HRP (A10260; Thermo-Fisher Scientific), and α-Myc-HRP (SAB4200742; Sigma-Aldrich).

### Kiwifruit transformation

*A. tumefaciens*-mediated transformation of *A. chinensis* was performed as previously described ^40^. The medium used for kiwifruit transformation in this study was adapted from Wang *et al*. ^40^ (Table S3). Briefly, leaf strips excised from *in vitro*-grown shoots were inoculated with suspension culture of *Agrobacterium* strain EHA105 (at O.D∼0.8–1.0) with infection buffer (M2) for 45 min. Inoculated leaf strips were transferred to co-cultivation medium (M3) and incubated at 24°C for 2 days. After co-cultivation, the leaf strips were transferred to regeneration and selection medium containing kanamycin 150 mg/L (M4). Individual calli were excised from the leaf strips for further selection and bud induction, and adventitious buds regenerated from the calli were excised and transferred to shoot elongation medium (M5). When shoots had grown to 1–2 cm high, they were transplanted onto rooting medium (M6).

### *In planta* bacterial growth quantification assays

Resistance to Psa was determined by flooding tissue culture-grown plantlets or leaf-dip for glass house-grown plants with Psa inoculum diluted to 10^7^ cfu/mL in 10mM MgCl_2_ supplemented with 0.025% Silwet™ L-77 surfactant (PhytoTechnology Laboratories®). Tissue culture plantlets were flooded for 3 min each. For glasshouse plants, leaves were submerged in inoculum for 20 s. Leaves were sampled in quadruplicate at selected time points using a using a 1-cm^2^ cork borer with each replicate carrying four leaf discs, ground in 10mM MgSO_4_ using a Storm24 Bullet Blender (Next Advance, United States). For tissue culture experiments, Psa growth was quantified by plating a 10-fold dilution series onto LB agar. Samples were incubated for 2 days before colony counting to calculate cfu/mL.

For both tissue culture and glasshouse experiments, Psa was also quantified using a qPCR method adapted from that of Hemara *et al.* ^28^. Briefly, genomic DNA was extracted from leaf samples using the PDQeX nucleic acid extractor (MicroGEM, New Zealand). Real-time qPCR was carried out on an Illumina Eco real-time PCR platform using SsoFast EvaGreen Supermix (BioRad). qPCR was performed using Psa intergenic transcribed spacer (*ITS*) and plant *AcEF1α* primers (Table S2). Relative Psa biomass was calculated by normalising the cycle threshold (Ct) values for Psa *ITS* to those of the kiwifruit *AcEF1α* reference gene and plotting the Ct difference for each sample. Results were plotted in R using the ggplot2 and ggpubr packages, with significance values calculated using a two-tailed Welch’s post hoc t-test.

### Disease phenotyping assays

Plantlets were flood-inoculated with Psa inoculum at approximately 10^6^ cfu/mL. Individual plantlets were scored weekly on a scale of 0 (asymptomatic) to 4 (dead) for 6 weeks post-infection (dpi). Disease phenotypes were quantified using an area under the disease progression curve (AUDPC) adapted from previous work (Schandry 2017). Briefly, mean disease index (DI) scores were plotted for weeks 2–6. AUDPC values were calculated for each sample and fitted to a linear mixed effects model using the lme4 package ^44^. Post hoc Tukey’s tests were performed via multcomp package ^45^. All statistical analyses were conducted in R, with figures produced using the ggplot2 package ^46,47^.

### Biolistic co-bombardment reporter eclipse assay

The biolistic report eclipse assay was described previously ^33^. Six bombardments were performed for each effector in an experiment, and carried out in triplicate (technical replicates) and expressed in terms of ρmol of MU produced per minute per gram of total protein. The means and standard errors were calculated from the six replicates conducted per experiment, from three independent experimental runs (n = 18). Data for each treatment were stacked from all three runs and were analysed by ANOVA followed by a Tukey’s HSD post hoc test.

### Phylogenetic analyses

The sequence information of NbNRG1 (GenBank: DQ054580.1), AtRPS2 (GenBank: AF487807.1), AtRPM1 (GenBank: KC211321.1), MdMr5 (GenBank: KT013245.1), SlPtr1 (GenBank: MT134103.1), GmRPG1r (GenBank: KF958751.1), GmRPG1b (GenBank: AY452685.1), AtZAR1 (GenBank: AK227017.1), and NbZAR1

(GenBank: MH532570.1) were obtained from NCBI. The top four BLASTP hits of each AcPTR1 orthologues NB-ARC domains with at least 20% identical residues in the domain were identified in the *Nicotiana benthamiana* (version 0.4.4) genome sequence ^29^. In the phylogenetic tree, the NB-ARC domain of each protein hit was determined using Interpro scan (https://www.ebi.ac.uk/interpro/), and the nucleotide sequences of the NB-ARC domain for each contracts were aligned to check for the presence of the NB-ARC conserved motifs. Multiple nucleotide sequence alignment were created by using MUSCLE alignment in Geneious Prime. The nucleotide distance was calculated by GAMMA GTR model with the algorithm of rapid bootstrapping and search for best-scoring ML tree and phylogenetic trees were generated using RAxML method ^48^ with 100 bootstrap replicates in Geneious Prime.

## Acknowledgements

We thank Monica Dragulescu and G Wadasinghe for maintaining *N. benthamiana* plants in the glasshouse. Finally we thank Dr Joanna Bowen and Dr Erika Varkonyi-Gasic for critically reviewing the manuscript. This work was supported by Plant & Food Research’s kiwifruit royalty investment programme (KRIP).

## Contributions

CB, JJ, and SMY conceived the project and designed the experiments. SMY performed *Agrobacterium*-mediated transient expression experiments and analysed the data; MY conducted and analysed the co-IP experiments; SMY and TW performed kiwifruit transformation and maintained plantlet materials; RKYC performed and analysed biolistic transformation reporter eclipse assay; AC and KT performed disease phenotyping assays with plantlets; SS and AC performed and analysed *in planta* bacterial growth quantification assays; SS, LMH, and JJ assisted in statistical analysis and figure generation; SMY wrote the manuscript; CB, EHAR, JJ, and SS revised the manuscript.

---

**Table S1.**
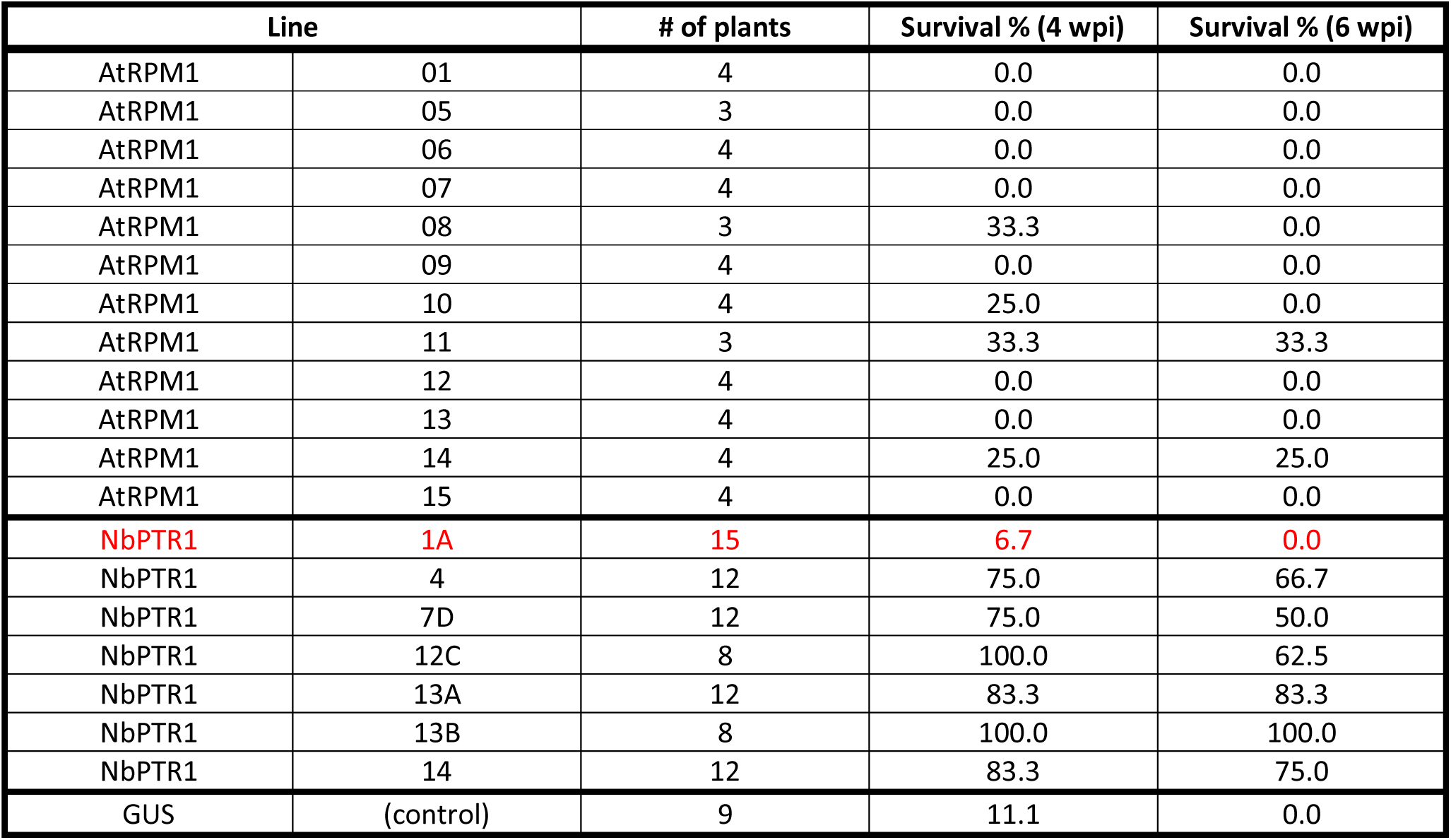
Comparison of survival rate at 4 and 6 weeks post-inoculation (wpi) between At*RPM1* kiwifruit transgenic plants, Nb*PTR1* transgenic plants, and *β-glucuronidase (GUS)* control transgenic plants, by flooding with Psa3 ICMP 18884 at 10^7^ cfu/mL. The very low NbPTR1 gene expression line 1A, is highlighted in red text. All genes (At*RPM1*, Nb*PTR1a*, or *GUS*) were cloned under the control of a 35S CaMV promoter.

| Line |  | # of plants | Survival % (4 wpi) | Survival % (6 wpi) |
| --- | --- | --- | --- | --- |
| AtRPM1 | 01 | 4 | 0.0 | 0.0 |
| AtRPM1 | 05 | 3 | 0.0 | 0.0 |
| AtRPM1 | 06 | 4 | 0.0 | 0.0 |
| AtRPM1 | 07 | 4 | 0.0 | 0.0 |
| AtRPM1 | 08 | 3 | 33.3 | 0.0 |
| AtRPM1 | 09 | 4 | 0.0 | 0.0 |
| AtRPM1 | 10 | 4 | 25.0 | 0.0 |
| AtRPM1 | 11 | 3 | 33.3 | 33.3 |
| AtRPM1 | 12 | 4 | 0.0 | 0.0 |
| AtRPM1 | 13 | 4 | 0.0 | 0.0 |
| AtRPM1 | 14 | 4 | 25.0 | 25.0 |
| AtRPM1 | 15 | 4 | 0.0 | 0.0 |
| NbPTR1 | 1A | 15 | 6.7 | 0.0 |
| NbPTR1 | 4 | 12 | 75.0 | 66.7 |
| NbPTR1 | 7D | 12 | 75.0 | 50.0 |
| NbPTR1 | 12C | 8 | 100.0 | 62.5 |
| NbPTR1 | 13A | 12 | 83.3 | 83.3 |
| NbPTR1 | 13B | 8 | 100.0 | 100.0 |
| NbPTR1 | 14 | 12 | 83.3 | 75.0 |
| GUS | (control) | 9 | 11.1 | 0.0 |

**Table S2.**
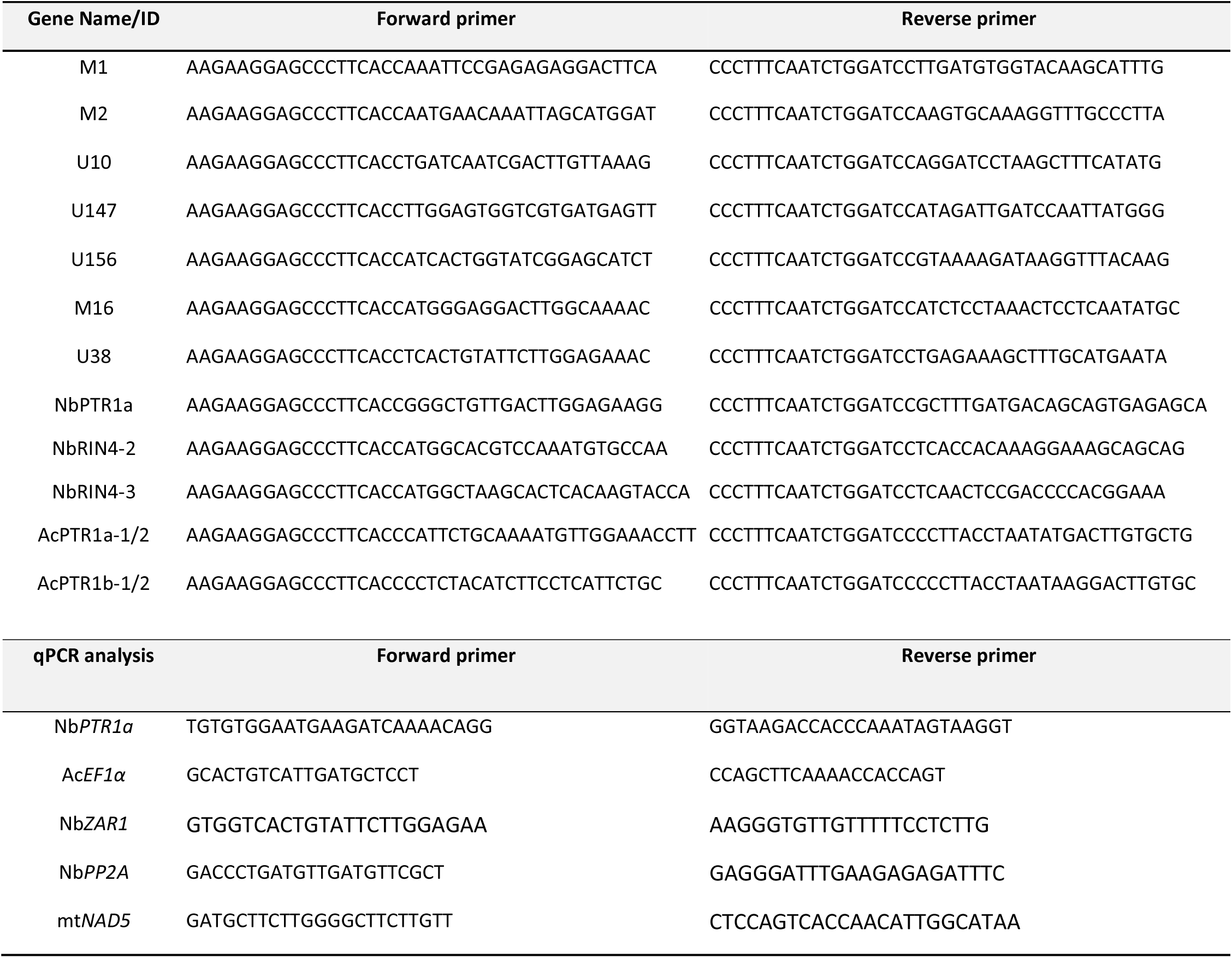
PCR Primers used in this study.

**Table S3.** Media used for kiwifruit transformation in this study.

| Medium | Ingredient |
| --- | --- |
| M1 | MS + indolebutyric acid (IBA) 0.2 mg/L + zeatin 0.2 mg/L+ sucrose 2% + agar (gemantown) 0.65%, pH: 5.8 |
| M2 | ½ MS + 100 uM acetosyringone + 2% sucrose, pH: 6.0 |
| M3 | ½ MS + zeatin 3.0 mg/L + 6-benzylaminopurine (BAP) 1.0 mg/L + 1-naphthaleneacetic acid (NAA) 0.1 mg/L + sucrose 3% + Phytigel 0.25% + acetosyringone 50uM, pH: 5.8. |
| M4 | MS + zeatin 3.0 mg/L + BAP 1.0 mg/L + NAA 0.1 mg/L + sucrose 3% + Phytigel 0.25% + kanamycin 150 mg/L + timentin 500 mg/L, pH: 5.8. |
| M5 | MS + zeatin 0.2 mg/L + sucrose 2% + agar (gemantown) 0.7% + kanamycin 50 mg/L + timentin 500 mg/L, pH: 5.8. |
| M6 | ½ MS + IBA 0.6 mg/L + sucrose 2% + agar 0.7%, pH: 5.8 |

## Supplementary experimental procedures

### RT-qPCR quantification of Nb*PTR1a*/Nb*ZAR1* in *N. benthamiana*

To check the expression of Nb*PTR1a/* Nb*ZAR1* with or without transient expression of silencing constructs in *N. benthamiana*, leaves were agroinfiltrated with silencing constructs m1 (anti-Nb*PTR1a*), m16+u38 (anti-Nb*ZAR1*), or GGT control vector, each at OD_600_ of 0.2. Leaf disks were collected at 2 days post-infiltration. RNA from leaf discs was extracted using the Spectrum Plant Total RNA kit (Sigma-Aldrich) and cDNA was synthesised using extracted RNA as the template and the QuantiTect Reverse Transcription kit (Qiagen) following the manufacturers’ instructions. Real-time qPCR was carried out on an Illumina Eco real-time PCR platform using SsoFast EvaGreen Supermix (BioRad). qPCR was performed using Nb*PTR1a*, Nb*ZAR1*, *N. benthamiana PROTEIN PHOSPHATASE 2A* (Nb*PP2A*)^49^, and plant-mitochondrial (*mtNAD5*) ^50^ primers (Table S2). Relative Psa biomass was calculated by normalising the cycle threshold (Ct) values for Nb*PTR1a* or Nb*ZAR1* to those of the Nb*PP2A* and *mtNAD5* reference genes and plotting the Ct difference for each sample. Results were plotted with significance values calculated using a two-tailed Student’s post hoc t-test.

### Disease phenotyping scoring

For tissue culture experiments, phenotypic scoring was carried out weekly for 6 weeks post-infection. Individual plantlets were scored from a scale of 0 to 4 (0: heathy; 1: mild symptoms; 2: medium symptoms: 3: severe symptoms; and 4: dead). There were several symptoms which were considered, such as necrosis/leaf wilting, leaf spots and white spots. Depending on the overall symptom extent, a scoring from 1 to 3 was given, where a score of 1 was given when any of these symptoms had just been observed or were beginning to appear, and gradually as the severity of these symptoms increased in the following weeks, a progressively larger score of 2 or 3 was given.

### Electrolyte leakage quantification in *A. chinensis* var. *chinensis* ‘Hort16A’

For electrolyte leakage assays in *A. chinensis* var. *chinensis* ‘Hort16A’ wild-type and transgenic plants, a similar approach was adapted from that of Jayaraman *et al*. ^33^. Overnight bacterial growth harvested directly from LB plates was re-suspended in 10mM MgCl_2_ and diluted to and inoculum of approximately 10^8^ cfu/mL (OD_600_ = 0.25). Young leaves (or 0.8-cm leaf discs) were harvested from tissue culture-grown kiwifruit plantlets before submersion in 20 mL of bacterial inoculum and vacuum infiltration. Infiltration was performed using a glass bell and a Rocker 300 oil-free vacuum pump (Rocker, Taiwan) with two 1-min bursts at 700 mm Hg. Leaf discs were sampled (for well-infiltrated leaves) and washed thoroughly before storing in 1.5 mL sterile MilliQ® water. Conductance readings were taken using a LAQUAtwin EC-33 conductivity meter (Horiba) at 0, 24, 48 and 72 hours post-infiltration. Means and standard errors were calculated from four biological replicates per sample.

---

**Figure S1.**
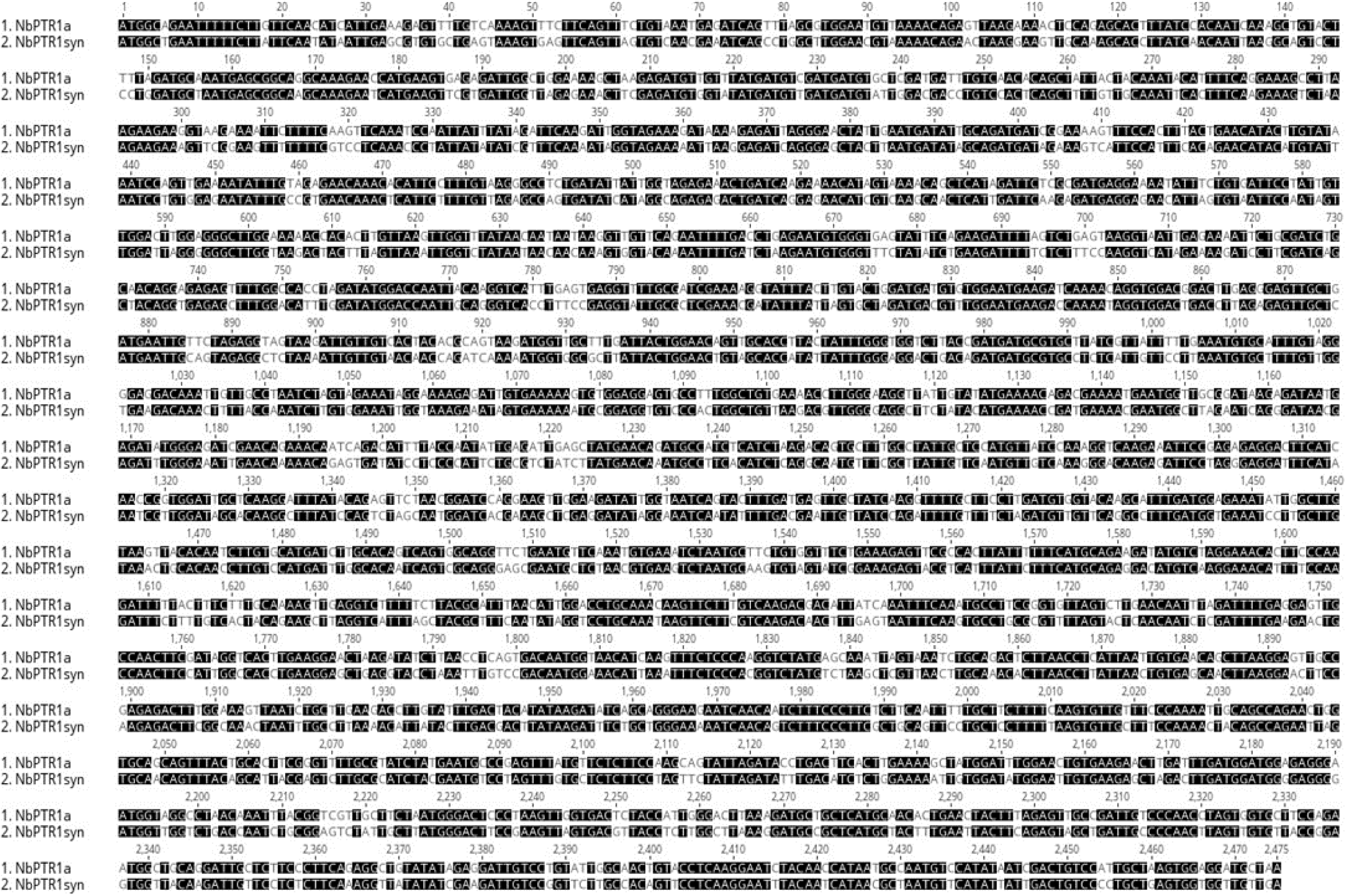
ClustalW alignment of wild-type Nb*PTR1a* and synthetic Nb*PTR1a* (Nb*PTR1syn*). Identical nucleotides are shown with black shading. The amino acid sequence of NbPTR1a is identical to that of NbPtr1syn.

**Figure S2.**
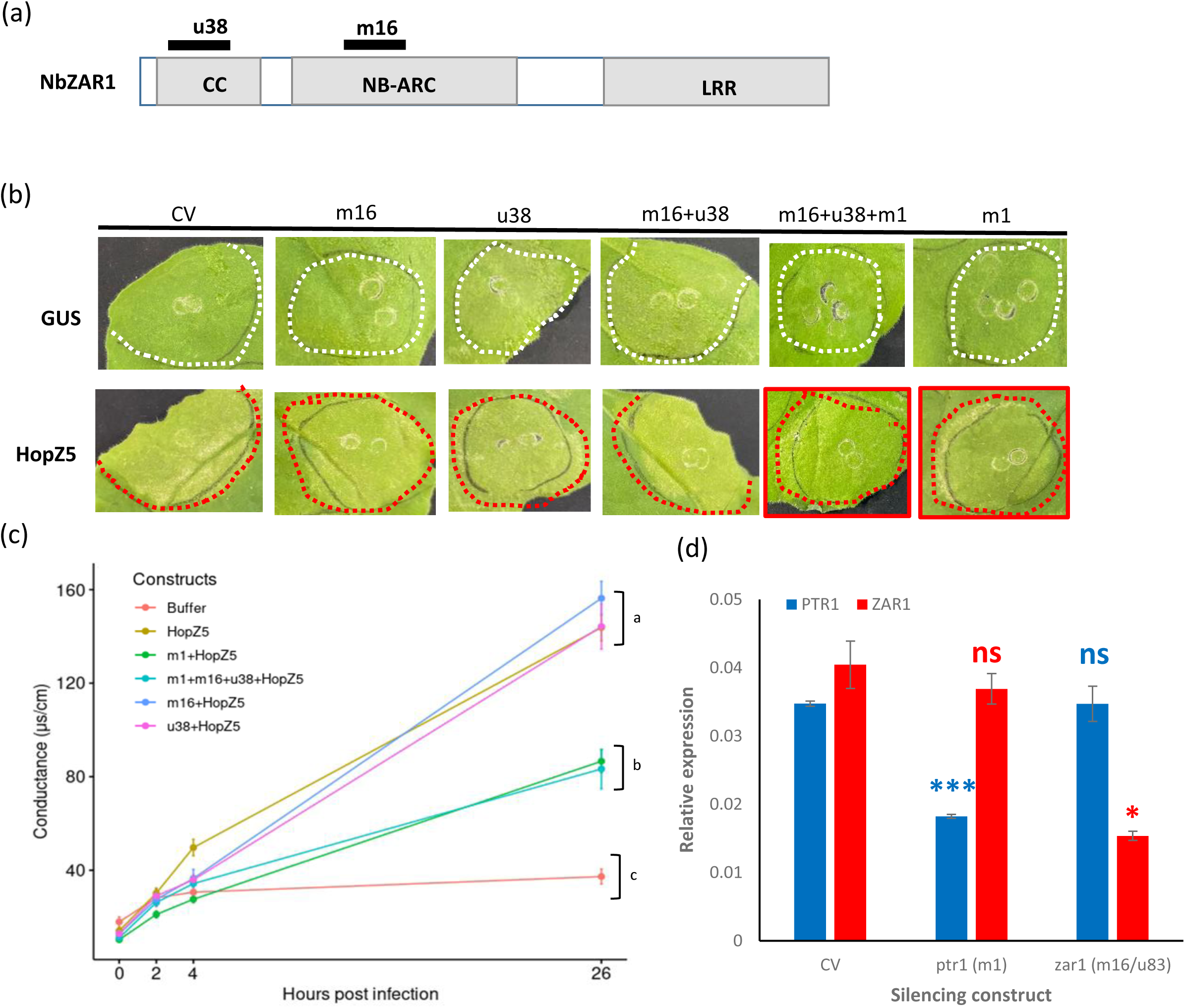
Two silencing fragments targeting Nb*ZAR1*, could not reduce HopZ5-triggered hypersensitive response in *Nicotiana benthamiana*. **a** Schematic of the NbZAR1 protein structure indicating the coiled-coil (CC), nucleotide binding site present in APAF-1, R proteins and CED-4 (NB-ARC), and leucine-rich repeat (LRR) domains. The position of the m16 fragment (16-136bp) and the u38 fragment (555-703bp) are indicated with black bars. **b** Silencing construct m1, but not m16 and/or u38, suppress HopZ5-triggered cell death. *N. benthamiana* leaves were agroinfiltrated with silencing constructs or GGT control vector (CV) at OD_600_ of 0.2 each, followed by agroinfiltration with HopZ5 or GUS (each at OD_600_ of 0.2) 48 hr later. Red dashed line indicates the HopZ5-infiltrated area and white dashed line for GUS-infiltrated area. Panels with reduced cell death in the HopZ5-infiltrated patches are marked with red borders. Leaves were photographed 4 days post-infiltration(dpi). The results shown are representative of at least 14 out of 20 independent leaves from different plants. **c** Conductivity of HopZ5-induced cell death with silencing constructs, m16 and u38, in *N. benthamiana*. Leaves were agroinfiltrated with silencing constructs m1, m16, u38, m16+u38, m1+m16+u38, or GGT control vector (CV), each at OD_600_ of 0.2, followed by agroinfiltration with HopZ5 or GUS (each at OD_600_ of 0.4) 48 h later. Leaf disks were collected at 1 day post final infiltration. Error bars represent the standard errors of the means for ten independent biological replicates, collected from two independent experimental runs (n=10). HopZ5 alone was used as positive control and infiltration buffer (10mM MgCl_2_, 100 μM acetosyringone) as a negative control. Letters indicate statistically significant differences from a one-way ANOVA and Tukey’s HSD *post hoc* test for values at 26 h post sampling. **d** qPCR quantification of Nb*PTR1a* or Nb*ZAR1* expression with silencing constructs, m1, m16 and u38, in *Nicotiana benthamiana*. Leaves were agroinfiltrated with silencing constructs m1, m16+u38, or GGT control vector (CV), each at OD_600_ of 0.2. Leaf disks were collected at 2 day post infiltration. The Nb*PTR1a* or Nb*ZAR1* expression is normalised to two reference genes, Nb*PP2A* and *mtNAD5*. Data are shown as means ± SEM of three biological replicates. Asterisks indicate results of a two-tailed Student’s t-test between the silenced sample and control vector (CV); (*) P < 0.05, (***) P < 0.001, and (ns) non-significant.

**Figure S3.**
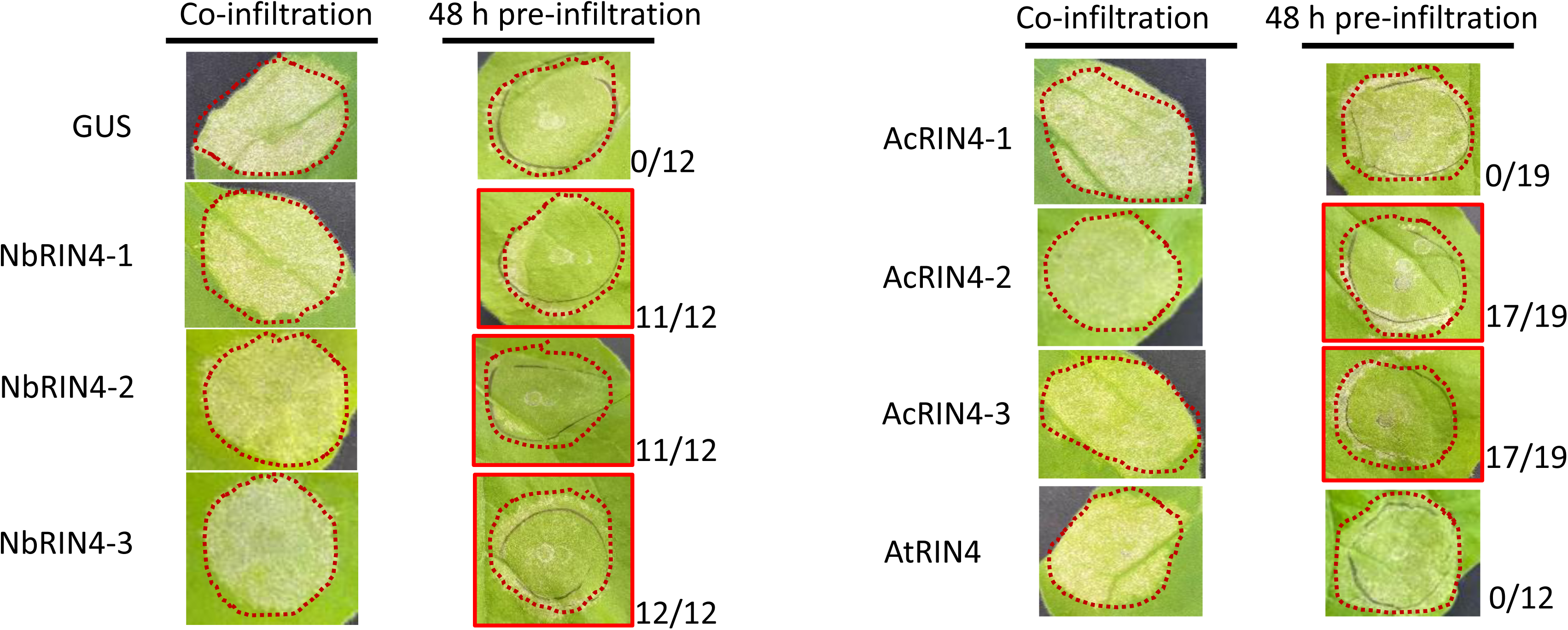
NbPTR1a-triggered autoimmunity can be reduced by either NbRIN4s or AcRIN4s. Transient expression of Nb*PTR1a* and Nb*RIN4* constructs, Ac*RIN4* constructs or At*RIN4*. Co-infiltration: *Nicotiana benthamiana* leaves were agroinfiltrated by Nb*PTR1a* (OD_600_ of 0.2) and GUS, Nb*RIN4*s, Ac*RIN4*s, or At*RIN4* (each at OD_600_ of 0.4); 48 h pre-infiltration: leaves were agroinfiltrated by GUS, Nb*RIN4*s, Ac*RIN4*s, or At*RIN4* (each at OD_600_ of 0.4), followed by agroinfiltration with Nb*PTR1a* (OD_600_ of 0.2) 48 h later. Leaves were photographed 4 or 5 days post-Nb*PTR1a* infiltration. Red dashed lines indicate the Nb*PTR1a*-infiltrated area. Fractions beside each panel indicate the incidence of patches with reduced cell death from pre-expression of RIN4. Red borders around panels indicate infiltrations where cell death counts were reduced in greater than 50% of infiltrations.

**Figure S4.**
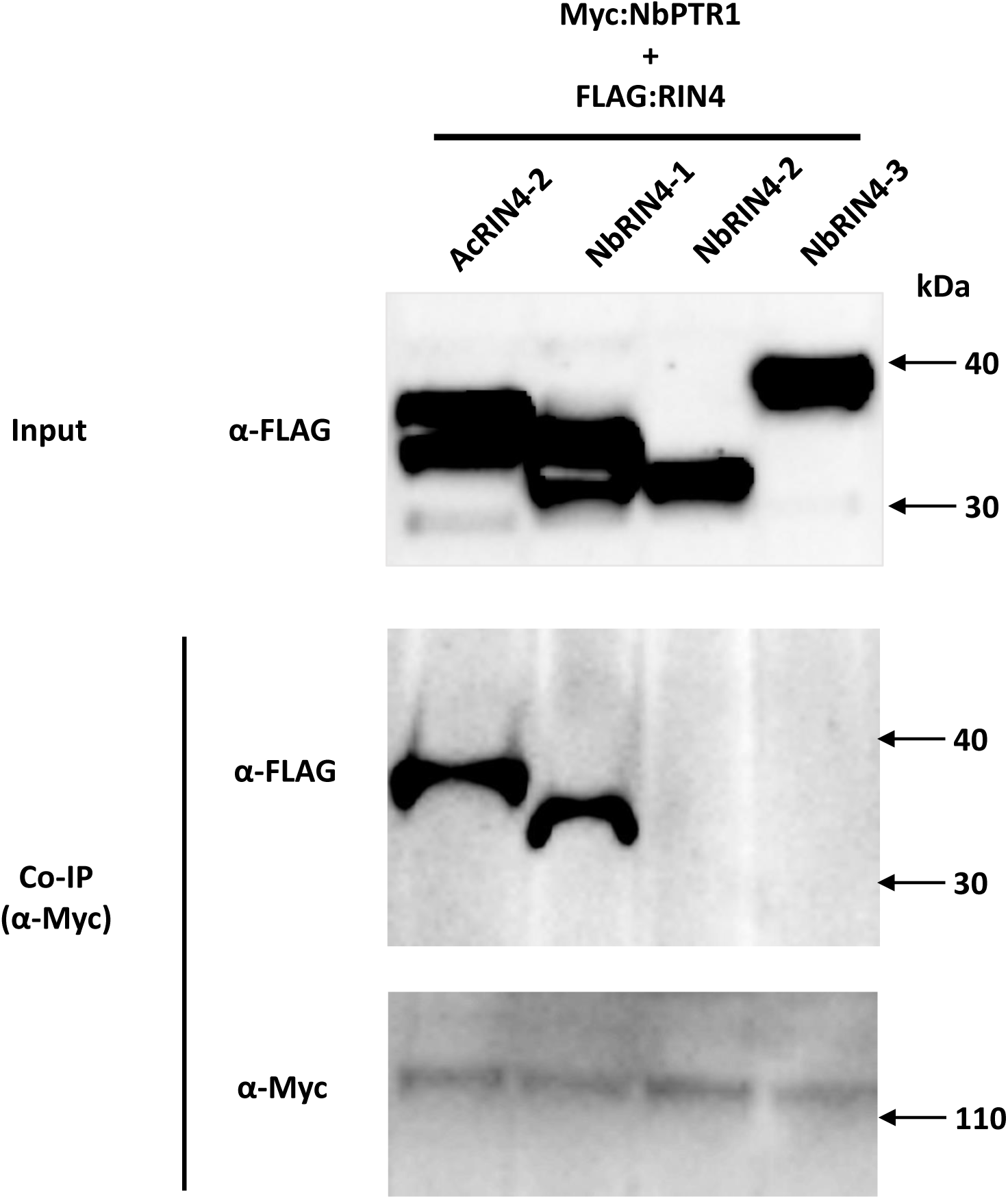
NbRIN4-1 stably interacts with NbPTR1a *in planta*. Co-immunoprecipitation of NbPTR1a and NbRIN4 homologues. NbPTR1a and NbRIN4 homologues were co-expressed simultaneously via agroinfiltration, each at OD_600_ of 0.1. 2 days post-infiltration, leaf samples were harvested, protein extracts prepared and precipitated using anti-Myc antibody. Western blot of input and precipitated proteins were probed with anti-FLAG or anti-Myc antibody. IP, immunoprecipitation.

**Figure S5.**
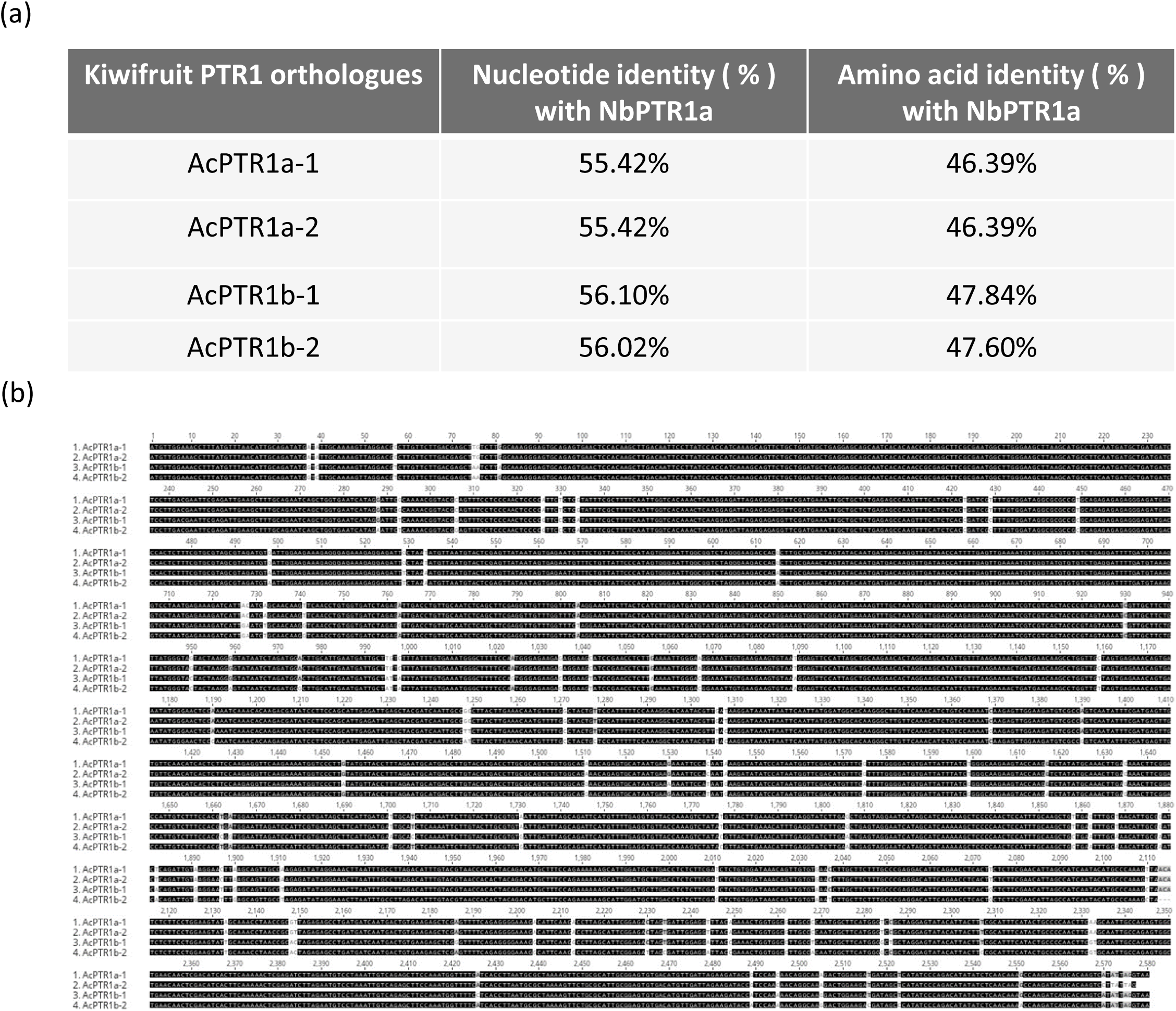
*PTR1a* orthologues in kiwifruit. **a** Comparison of Nb*PTR1a* nucleotide sequences with its orthologues in kiwifruit (Ac*PTR1a*). **b** Nucleotide sequence alignment of the four Ac*PTR1a*/Ac*PTR1b* orthologues cloned from Hort16A. Identical nucleotides are shown with black shading.

**Figure S6.**
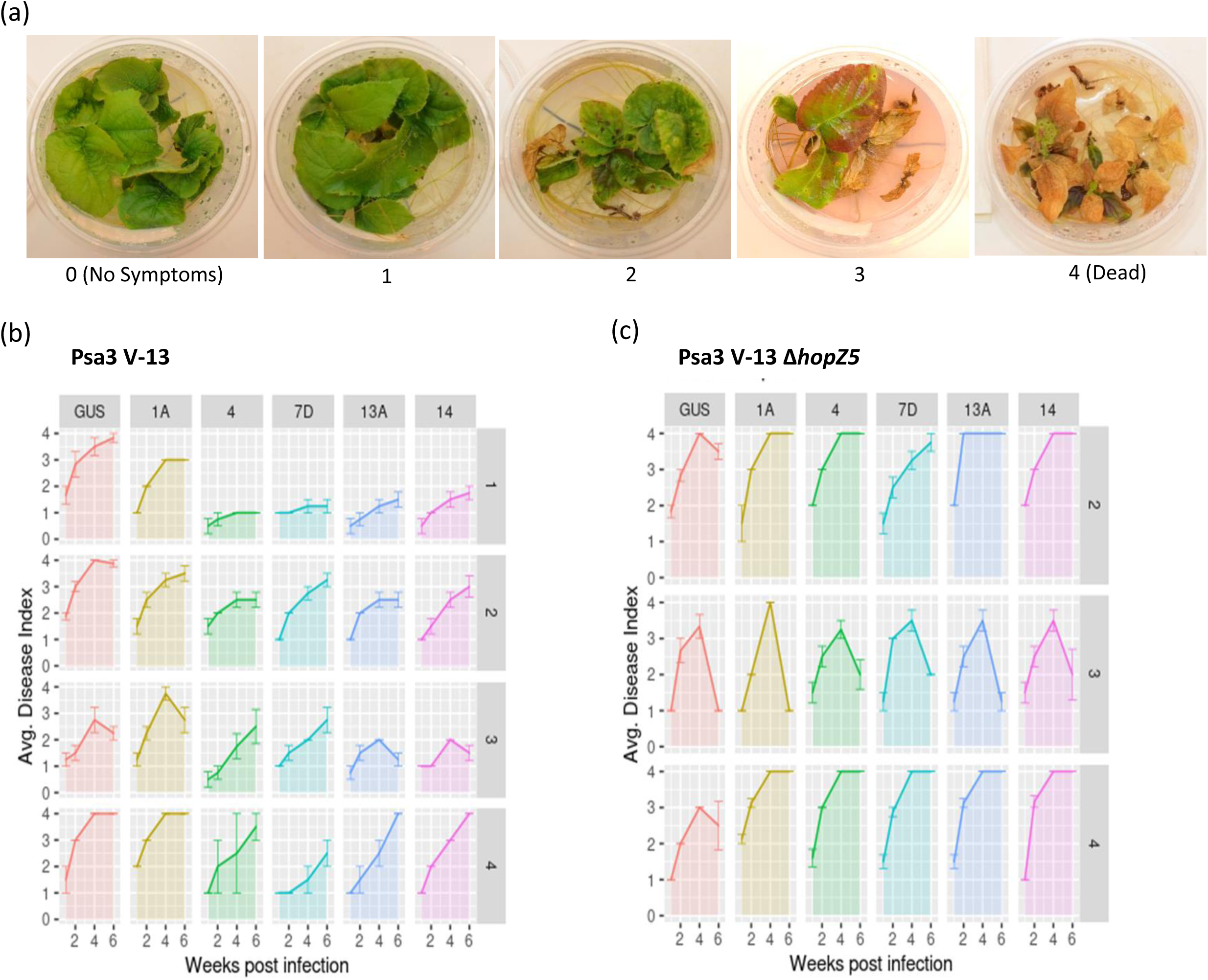
Disease phenotyping analysis of Nb*PTR1a* transgenic kiwifruit plants. **a** The scale of symptom development in Nb*PTR1a* kiwifruit transgenic plants in *Actinidia chinensis var. chinensis* ‘Hort16A’ post-inoculation with either Psa3 V-13 or Psa3 V-13 *ΔhopZ5*. Psa3 V-13 or Psa3 V-13 *ΔhopZ5* was flood-inoculated at approximately 10^6^ CFU/mL. Phenotypic scoring was performed every week until 6 weeks post-inoculation (wpi), where each plantlet was given an individual score from a scale of 0 (no symptoms) to 4 is (dead). **b, c** Disease area analysis of symptom scoring post inoculation (dpi) with either Psa3 V-13 (**b**) or Psa3 V-13 Δ*hopZ5* (**c**). Psa3 V-13 was flood-inoculated as described in (**a**), and the symptom scoring was monitored up to 6 wpi. This experiment was conducted four times with similar results (indicated on the right-hand y-axis labels; run 1 with Psa3 V-13 alone, runs 2-4 with both Psa3 V-13 and Psa3 V-13 *ΔhopZ5*).

**Figure S7.**
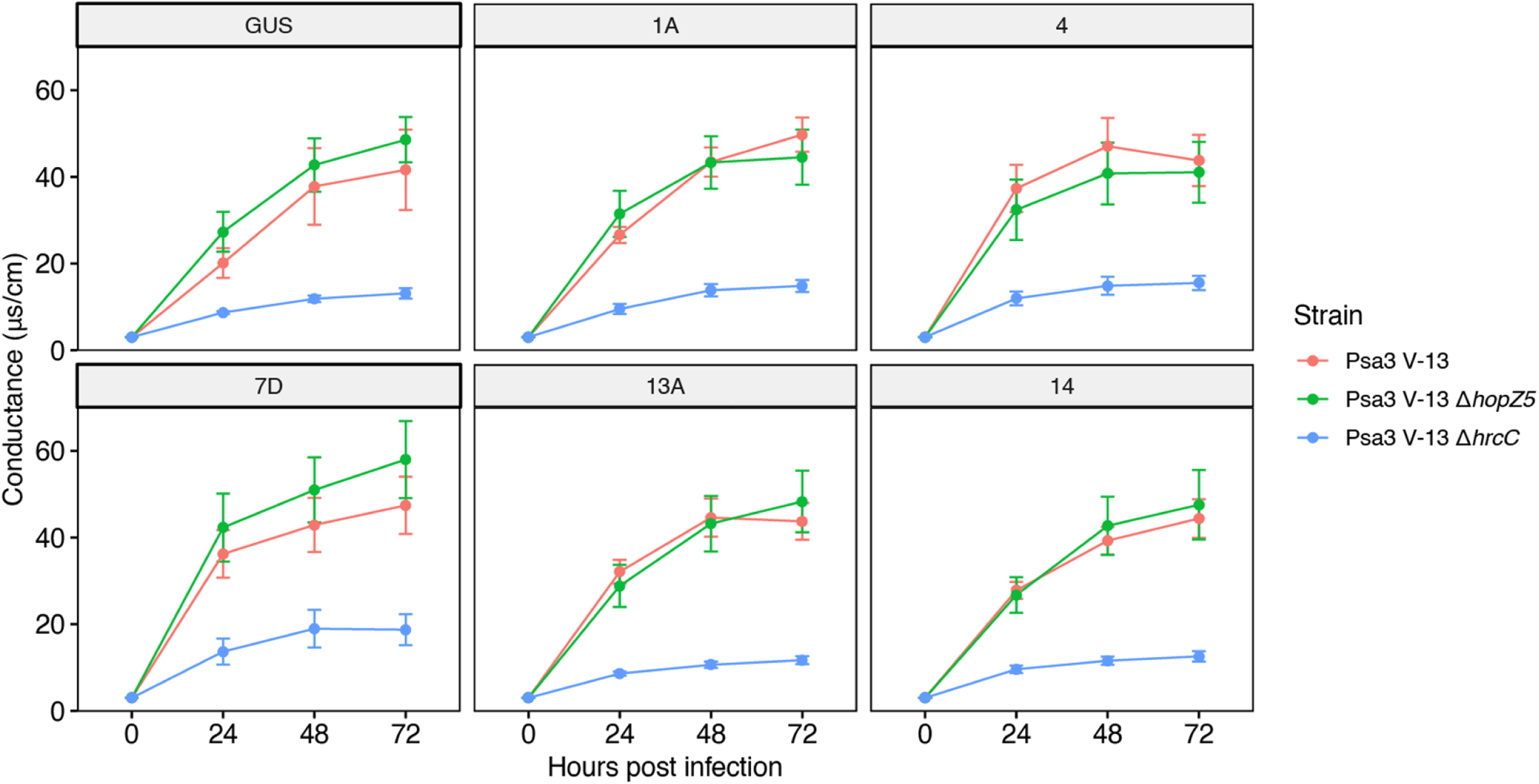
Psa V-13 does not trigger HopZ5-specific ion leakage in transgenic Nb*PTR1a* kiwifruit plants. Measurement of hypersensitive response (HR) of transgenic Nb*PTR1a* glasshouse-grown plants by electrolyte leakage. Leaf discs from glasshouse-grown plants were vacuum-infiltrated with Psa3 V-13, Psa3 V-13 *ΔhopZ5*, or Psa V-13 *ΔhrcC* inoculum at approximately 10^8^ CFU/mL, and electrical conductivity due to HR-associated electrolyte leakage was measured at selected time points over 72 h. Error bars represent the standard errors of the means calculated from the three independent experimental runs, each with five biological replicates.

**Figure S8.**
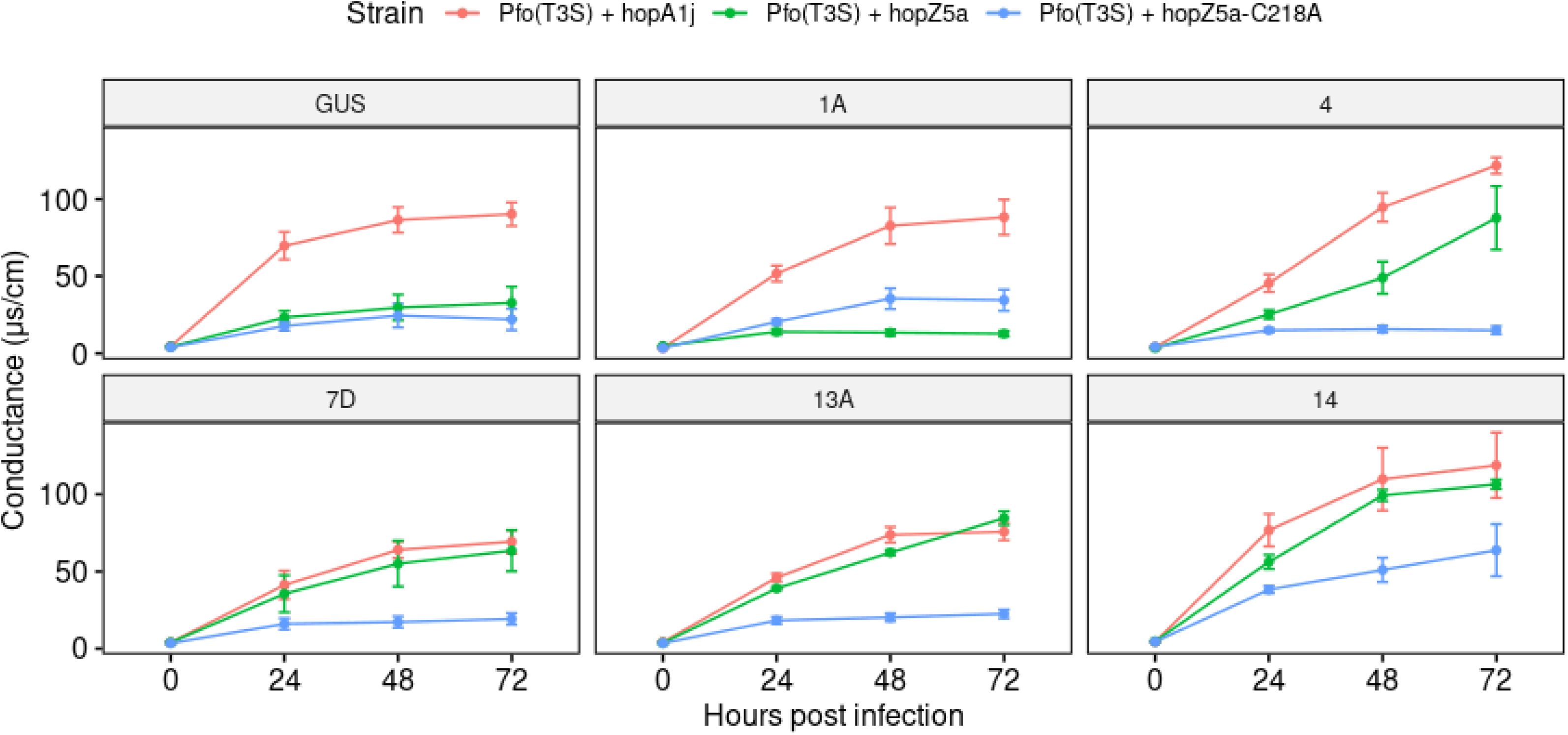
HopZ5 recognition in transgenic Nb*PTR1a* kiwifruit plants triggers weak ion leakage. Measurement of hypersensitive response (HR) of Nb*PTR1a* transgenic tissue culture-grown plantlets by electrolyte leakage. Leaf discs from plantlets were vacuum-infiltrated with Pseudomonas fluorescens PF0-1 (T3S) carrying positive control *hopA1j* (from *P. syringae* pv. *syringae* 61), *hopZ5*, or *hopZ5-*C218A (enzymatic dead mutant) inoculum at ∼10^8^ CFU/mL, and electrical conductivity due to HR-associated electrolyte leakage was measured at selected time points over 72 h. Error bars represent the standard errors of the means calculated from four biological replicates.

